# α-1,3-Glucan-Driven Remodeling of the Conidial Cell Wall in an *Aspergillus fumigatus* Vaccine Strain Alters Innate Immune Recognition

**DOI:** 10.64898/2025.12.31.697239

**Authors:** Kalpana Singh, Ankur Ankur, Jayasubba Reddy Yarava, Caroline Mota Fernandes, Gianluca Vascelli, Alessia Sulla, Teresa Zelante, Maurizio Del Poeta, Tuo Wang

## Abstract

*Aspergillus fumigatus* is a major cause of invasive aspergillosis in immunocompromised patients, where current antifungal therapies are limited by toxicity, drug resistance, and lack of durable protection, and no vaccines are available. A mutant lacking the sterylglucosidase-encoding gene (*sglA*) has emerged as a candidate that induces protective immune responses, but the structural basis for this phenotype remains unclear. Here, we use cellular solid-state NMR spectroscopy to compare the organization of the conidial cell wall in Δ*sglA* and its wild-type counterpart. The Δ*sglA* conidial cell wall displays extensive remodeling, including increased α-1,3-glucan content and structural polymorphism, strengthened interactions with β-glucans, reduced hydration, and restricted molecular motion, together consolidating a more rigid scaffold with limited β-glucan accessibility. These structural changes are associated with altered neutrophil responses and a shift in innate immune signaling. This work links cell-wall reorganization to altered immune recognition in this vaccine candidate, with implications for future immunotherapeutic strategies.

**TEASER:** Molecular-level Insights from a fungal vaccine candidate show how cell-wall remodeling could affect immune response.

## INTRODUCTION

*Aspergillus fumigatus* is the major etiological agent of invasive aspergillosis, a life-threatening fungal infection that affects approximately 200,000 people worldwide each year and is associated with a mortality rate of 30-90%^1,2^. The disease primarily affects immunocompromised individuals, including patients with AIDS, organ or stem-cell transplants, and those receiving immunosuppressive therapies^2,3^. Infection is initiated by inhalation of airborne conidia, which reach the lung parenchyma and germinate into invasive hyphae^4–6^. In immunocompetent hosts, the innate immune response is generally sufficient to clear the fungus; however, in individuals with impaired immunity, the infection progresses to invasive disease^7–9^.

Azole antifungal drugs, which inhibit ergosterol biosynthesis, remain the first-line therapy for invasive aspergillosis^10–13^. Their clinical effectiveness, however, has been increasingly compromised by the emergence and global spread of azole-resistant *A. fumigatus* strains^14–16^. Echinocandins, a newer class of antifungals that inhibit β-1,3-glucan synthesis in the fungal cell wall, are also used in clinical practice^17–19^. Although effective in limiting fungal growth, echinocandins are largely fungistatic rather than fungicidal against *Aspergillus* species, and thus are primarily employed as second-line agents in invasive aspergillosis^20–22^. Compounding these therapeutic limitations, there is currently no antifungal vaccine licensed for the clinical prevention or treatment of invasive aspergillosis^23,24^. This challenge is further exacerbated by the fact that invasive aspergillosis primarily affects immunosuppressed patients, whose reduced immune-cell function compromises both natural host defenses and the effectiveness of potential vaccination strategies^24,25^.

In response to this unmet need, vaccination-based approaches have gained increasing attention. Several studies have demonstrated that vaccination with viable *Aspergillus* conidia can elicit protective immunity, with up to 70% of immunocompromised mice surviving lethal challenge following immunization^26,27^. Building on this concept, a gene-deletion strategy targeting sterylglucosidase (SGL1) in *Cryptococcus neoformans* generated a Δ*sgl1* mutant that accumulates sterylglucosides and confers complete protection against lethal fungal infections across multiple immunosuppression models^28,29^. A similar phenotype was observed in *A. fumigatus*, where the Δ*sglA* mutant exhibited attenuated virulence during primary infection and was fully cleared from the lungs of immunocompromised mice^30^. These advances suggest that fungal sterylglucosides, together with associated cell-wall alterations, may represent a promising immunomodulatory platform for vaccine development in vulnerable hosts.

However, the mechanisms underlying the avirulence of the vaccine-candidate strain, the *A. fumigatus* Δ*sglA* mutant, remain poorly understood, particularly with respect to how cell-wall remodeling and sterylglucoside accumulation contribute to its immunoprotective response. In this study, we employ ^13^C and ^1^H-detected solid-state NMR to uncover the unique structural features of the cell wall in intact Δ*sglA* conidia. This non-destructive spectroscopic approach enables the resolution of the structure, dynamics, and physical interactions of polysaccharides in living cells^31–33^. This capability has recently been leveraged to elucidate adaptive cell-wall remodeling across diverse fungal pathogens—including *Candida* species, *Cryptococcus* species, and filamentous fungi such as *Aspergillus* and *Mucor* species—in systems of major clinical and biological relevance^34–41^. High-resolution analyses of Δ*sglA* conidia reveal a distinct molecular architecture characterized by an unusually rigid and dehydrated cell wall, with markedly increased α-1,3-glucan incorporated into the rigid structural scaffold. We further correlate this α-1,3-glucan-mediated masking of β-glucans with the loss of Dectin-1 signaling in neutrophils, thereby bridging the gap between cell-wall remodeling and immune responses that underlie the functional mechanism of this vaccine candidate, and providing structural insights that may inform the future development of antifungal vaccines.

## RESULT

### Structural features of polysaccharides in the organization of *A. fumigatus* conidial cell walls

Prior to characterizing the vaccine-candidate strain, we first examined the polysaccharide structure and distribution in the wildtype control Δ*akuB*^KU80^, the parental strain with enhanced homologous recombination used to generate the Δ*sglA* vaccine mutant strain^42^. Intact, uniformly ^13^C-labeled conidia were analyzed directly by solid-state NMR, while scanning electron microscopy (SEM) confirmed that the cells retained the characteristic morphology of resting conidia and an intact cell-wall layer (**Figure 1A**). In the wild-type sample, the 2D ^13^C-^13^C CORD correlation^43^ spectrum acquired using dipolar-based ^1^H-^13^C cross-polarization (CP), which emphasizes rigid carbohydrates, was dominated by β-1,3-glucan signals, with weaker contributions from chitin (**Figure 1B** and **Fig. S1**). Minor signals from chitosan and β-1,6-glucan were also detected. These results reveal that the rigid core of the conidial cell wall is composed primarily of β-1,3-glucan, with additional support from chitin, chitosan, and β-1,6-glucan (**Figure 1C**). Three magnetically distinct chitin forms were identified based on their resolved C1-C2 cross-peaks (**Figure 1D**) and complete carbon connectivities (**Fig. S2**), reflecting local structural perturbations such as differences in conformation and hydrogen-bonding patterns.

**Figure 1.**
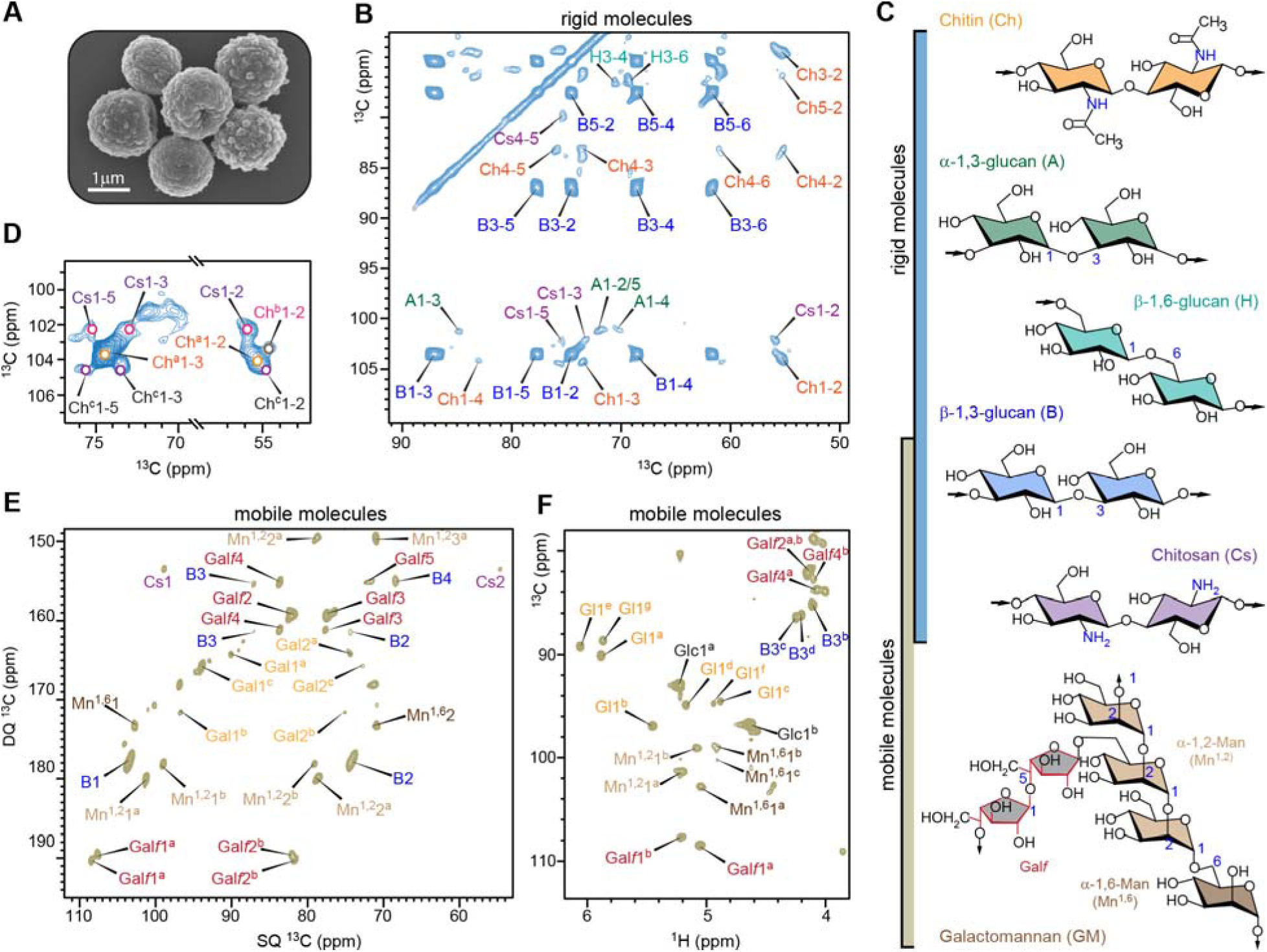
Rigid and mobile polysaccharides of wildtype *A. fumigatus* conidial cell wall. (**A**) Scanning electron microscopy (SEM) image of *A. fumigatus* dormant conidia (scale bar: 1 μm). (**B**) 2D ^13^C-^13^C correlation spectrum of *A. fumigatus* conidia acquired using a 53-ms CORD experiment, which selectively detects rigid molecular components through ^1^H-^13^C cross polarization (CP). Carbon resonance assignments for chitin (Ch), β-1,3-glucan (B), α-1,3-glucan (A), chitosan (Cs), and β-1,6-glucan (H) are shown in orange, blue, green, pink, and cyan, respectively. Each cross peak represents a correlation between two carbon atoms; for example, B1-3 corresponds to the correlation between C1 and C3 of β-1,3-glucan. (**C**) Summary of the NMR abbreviations used in this study, along with simplified structures of the major cell wall polysaccharides and their heterogeneous mobilities. (**D**) Expanded view of the 2D ^13^C-^13^C correlation spectrum highlighting a zoomed-in region that resolves three distinct chitin forms (Ch^a^, Ch^b^, and Ch^c^). (**E**) 2D ^13^C DP refocused *J*-INADEQUATE spectrum detecting mobile polysaccharides. Assignments contain NMR abbreviation and carbon number, for example, B1 represents β-1,3-glucan carbon 1. Glucofuranose: Gal*f*; α-1,2-mannose: Mn^1,2^; α-1,6-mannose: Mn^1,6^; galactose units: Gal. (**F**) Selected carbohydrate regions from the 2D hcCH TOCSY (DIPSI-3) spectrum of A. fumigatus conidia, showing signals from galactofuranose (Gal*f*), mannose units, glucans, and chitosan, as well as small molecules such as glucose (Glc) and galactose- or glucose-derived species (Gl).

Mobile polysaccharides were identified using a combination of ^13^C direct polarization (DP) and a short recycle delay in the 2D refocused J-INADEQUATE experiment^44^, which selectively suppresses signals from rigid components with slow ^13^C spin-lattice relaxation. The resulting spectrum showed well-resolved, sharp resonances from β-1,3-glucan, α-1,2-mannose, α-1,6-mannose, and galactofuranose (Gal*f*) units (**Figure 1E**). The latter three residues arise from galactomannan, whose backbone is composed of α-1,2- and α-1,6-linked mannose residues and is further substituted with β-1,5-linked (and sometimes β-1,6-linked) Gal*f* sidechains in *A. fumigatus* cell walls (**Figure 1C**).^45,46^. Thus, the mobile phase of the dormant conidial cell wall consists predominantly of β-1,3-glucan and galactomannan. The identification of β-1,3-glucan in two dynamically distinct domains reveals its dual structural role, extending from the rigid inner scaffold into the mobile outer matrix and thereby bridging rigid chitin with mobile galactomannan.

Structural polymorphism was also observed among the mobile carbohydrates. Two distinct forms of Gal*f* and two forms of α-1,2-linked mannose were resolved in both the ^13^C-detected spectrum (**Figure 1E**) and the ^1^H-detected 2D *J*-hcCH TOCSY spectrum (**Figure 1F**), with ¹H detection providing enhanced sensitivity for detailed structural analysis^47^. Notably, two Gal*f* forms were also detected in the mycelial cell wall, indicating that the galactomannan side chains exhibit similar structural complexity in both conidial and mycelial walls^48^. In addition, the 2D *J*-hcCH TOCSY spectrum^49^ resolved three forms of β-1,3-glucan and three forms of α-1,6-linked mannose, whose complete carbon connectivities and chemical shifts were further confirmed in an extended 3D *J*-hCCH TOCSY dataset (**Fig. S3**). This structural polymorphism reflects the molecular complexity of the soft matrix, in which polysaccharides may adopt diverse linkage and cross-linking patterns or sample multiple conformational energy minima.

### Deletion of *sglA* alters wall composition and increases α-glucan content and polymorphism

Compared with wildtype cells, the Δ*sglA* mutant displayed a slightly larger cell diameter, increasing from 1.8 µm to 1.9 µm (**Figure 2A** and **Table S1**). At the molecular level, we observed a pronounced change in the rigid polysaccharide composition: the content of α-1,3-glucan increased substantially in the mutant relative to the wild type (**Figure 2B**). This is evidenced by the enhanced α-1,3-glucan C1 and C3 peaks (A1 and A3) in the 1D ^13^C CP spectra (**Figure 2B**), as well as the appearance of strong intramolecular cross-peaks within α-1,3-glucan, such as A3-2/5, A3-4, and A3-6, in the 2D ^13^C-^13^C correlation spectra (**Figure 2C**). Peak-volume analysis indicated that the molar fraction of α-1,3-glucan in the rigid cell-wall core increased from 2% in the wild type to 20% in the mutant, accompanied by a decrease in β-1,3-glucan from 78% to 67% (**Figure 2D** and **Table S2**). This marked shift in glucan composition suggests a reorganization of the conidial cell wall upon deletion of *sglA* gene, with α-1,3-glucan providing structural integrity to the rigid cell wall, rather than a rigid matrix composed primarily of chitin/chitosan and β-glucan as in the wildtype.

**Figure 2.**
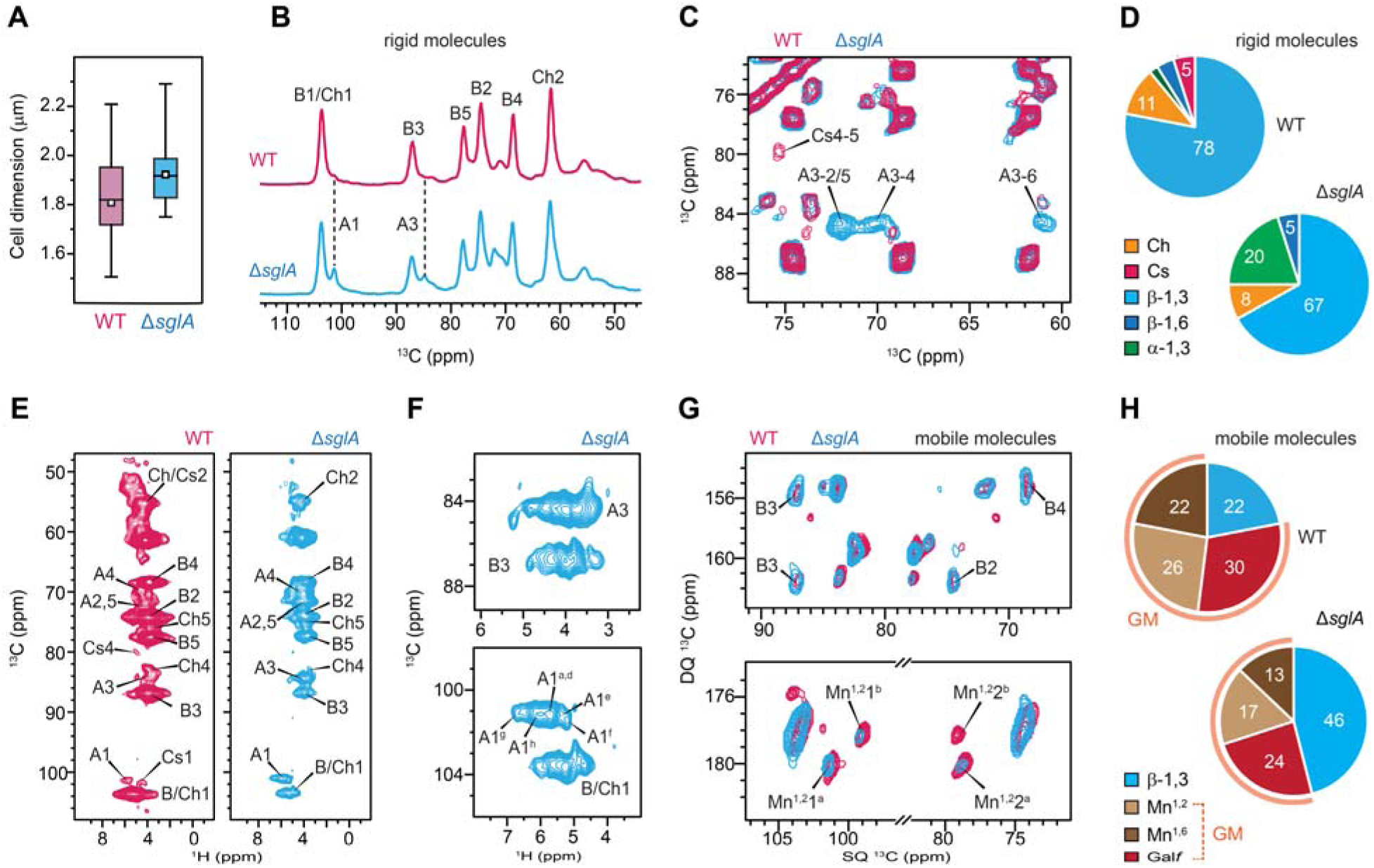
Enhancedα-glucan content, polymorphism, and altered composition in ΔsglA cell walls. (**A**) Cell diameter measured from SEM images. Boxes represent the interquartile range (IQR), with whiskers extending to 1.5 × IQR. Open squares indicate the mean, and horizontal lines indicate the median. Sample sizes: WT (n = 9) and Δ*sglA* (n = 9). Statistical analysis was performed using a t-test with one-tail comparison between WT and mutant strains. Statistically significant: *p-value[≤[0.05. (**B**) 1D ^13^C CP spectra showing rigid polysaccharides in *A. fumigatus* WT (top, magenta) and the Δ*sglA* mutant (bottom, blue). Dashed lines mark α-glucan peaks that emerge in the mutant. (**C**) 2D ^13^C-^13^C CORD correlation spectra showing signals from rigid polysaccharides in WT (blue) and Δ*sglA* (magenta). (**D**) Molar composition of rigid polysaccharides in WT and Δ*sglA*, estimated from peak volume analysis of the 2D CORD spectra. (**E**) Identification of rigid polysaccharides using a CP-based 2D hCH experiment with a short second CP contact time (50 μs) to detect one-bond ^13^C-^1^H correlations. (**F**) Zoomed region of the Δ*sglA* mutant spectrum showing distinct polymorphic forms of α-1,3-glucan. (**G**) Mobile molecules in WT (magenta) and Δ*sglA* (blue) detected by 2D ^13^C-DP refocused J-INADEQUATE spectra. (**H**) Molar composition of mobile polysaccharides in WT and Δ*sglA*, analyzed from peak volumes in the 2D DP J-INADEQUATE spectra. NMR abbreviations are as follows: B, β-1,3-glucan; Ch, chitin; chitosan, Cs; A, α-1,3-glucan; GM, galactomannan; Mn^1,2^, α-1,2-mannose; Mn^1,6^, α-1,6-mannose; Gal*f*, galactofuranose.

The ^1^H-detected hCH spectra revealed that the mutant is not only enriched in α-glucan, as indicated by the increased intensity of its carbon-1 peak (A1 in **Figure 2E**), but that this polymer also exhibits substantial structural polymorphism. This is evidenced by the broad distribution of signals for its ^1^H3 and ^1^H1 sites, with six magnetically nonequivalent ^1^H1 environments resolved for α-glucan (**Figure 2F**).

We also observed that the combined abundance of chitin and its deacetylated form, chitosan, decreased from 16% to 8% (**Figure 2D**). Notably, all chitosan, representing approximately 5% of the rigid molecules in the wildtype, was absent in the mutant, as indicated by the loss of the Cs4-5 cross-peak (**Figure 2C**) and the disappearance of all characteristic chitosan carbon signals (**Fig. S4**). Thus, in the Δ*sglA* mutant, chitin is not only reduced in amount but also remains fully acetylated, with no detectable chitosan.

Similar to the wildtype, the mutant retained a binary mobile phase composed of β-1,3-glucan and galactomannan; however, the peak intensity of galactomannan decreased substantially (**Figure 2G** and **Fig. S5**), resulting in a reduction of its molecular fraction in the mobile matrix from 78% to 54% (**Figure 2H** and **Table S3**). Together, these compositional and structural changes indicate a major remodeling of both the rigid core and the mobile matrix in the Δ*sglA* cell wall, characterized by a redistribution of a portion of β-glucan from the rigid core to the mobile phase, along with an expansion of α-1,3-glucan accompanied by reductions in chitin, chitosan, and galactomannan.

### Consolidated Δ*sglA* conidial cell wall with increased rigidity, dehydration, and dense packing

To probe sub-nanometer spatial proximities between polysaccharides, we performed 2D hChH experiments with RFDR-XY16 mixing on both wild-type and mutant samples (**Figure 3A**). This experiment produced additional intensities arising from long-range intra- and intermolecular correlations that were absent in the hCH spectrum, which primarily detects one-bond ^1^H-^13^C correlations. In the wild-type sample, only a few new long-range intramolecular cross-peaks were observed, including correlations between β-1,3-glucan carbon sites and its H1 proton (B4-B^H^1, B5-B^H^1, and B3-B^H^1), as well as between chitin C1 and its methyl proton (Ch1-^H^Me). In contrast, the Δ*sglA* mutant displayed clear intermolecular contacts (**Figure 3A, B**). Notable cross-peaks appeared between the H1 of α-1,3-glucan and multiple β-1,3-glucan carbon sites (B4-A^H^1, B2-A^H^1, B5-A^H^1, and B3-A^H^1), as well as between α-1,3-glucan C1 and the β-1,3-glucan H1 (A1-B^H^1). These interactions indicate that the abundant α-1,3-glucan uniquely observed in the mutant is effectively integrated with β-1,3-glucan on sub-nanoscale and has been effectively incorporated into the rigid core of the Δ*sglA* cell wall, providing additional structural reinforcement by acting as an adhesive molecule between various polysaccharides.

**Figure 3.**
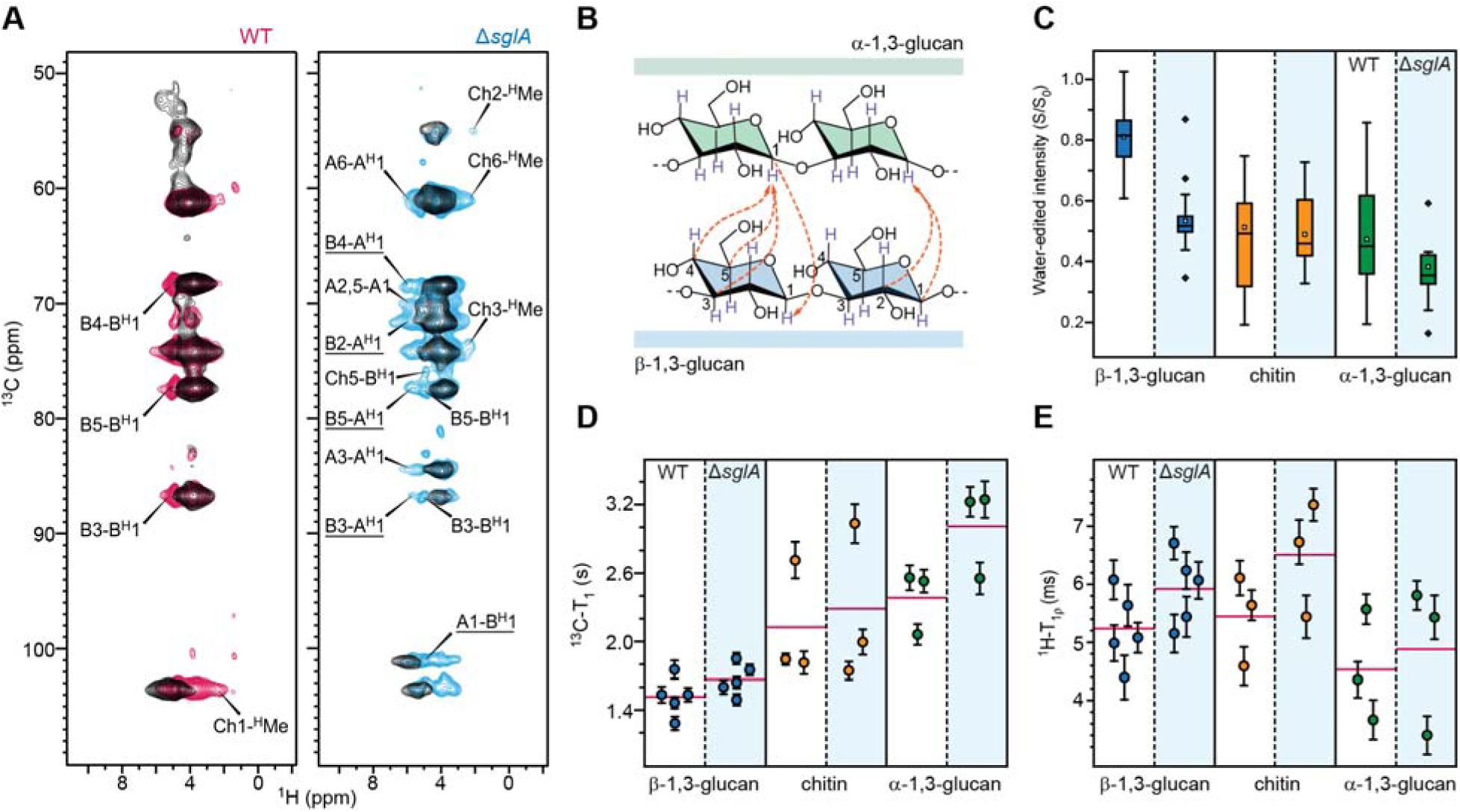
Hydration and dynamics of polysaccharides in the *A. fumigatus* conidial cell wall. (**A**) 2D hChH spectra (0.8 ms RFDR) of WT (magenta) and Δ*sglA* (blue) strains. One-bond 2D hCH spectra are overlaid in black for comparison. Intermolecular cross-peaks between α-1,3- and β-1,3-glucans are underlined. (**B**) Schematic summary of intermolecular glucan interactions detected in Δ*sglA* (orange dashed lines). Arrowheads indicate polarization-transfer direction. For example, a cross-peak at C3 of β-1,3-glucan arising from the ^1^H of α-1,3-glucan is labeled B3-A^H^1. (**C**) Box-and-whisker plots of water-edited intensities (S/S_0_) for β-1,3-glucan (blue; n=26, 30), α-1,3-glucan (green; n=15, 12), and chitin (orange; n=15, 9). Boxes show IQRs; whiskers extend to 1.5× IQR; outliers are stars. Means are open squares; medians are horizontal lines. (**D**) ^13^C-T_1_ relaxation times for β-1,3-glucan (blue; n=5, 5), α-1,3-glucan (green; n=3, 3), and chitin (orange; n=3, 3). (**E**) ^1^H-T_1_ relaxation times for the same polysaccharides. In (**D**-**E**), magenta lines show averages across carbon sites; error bars indicate s.d. of the fit parameters into a single-exponential equation.

With the introduction of a substantial amount of α-1,3-glucan and its tight integration into the rigid core, it is not surprising that the Δ*sglA* cell wall becomes more dehydrated than that of the wild type. This is reflected by the reduced water-edited intensities (S/S_0_), which report the extent of water association at individual carbohydrate sites (**Figure 3C**; **Table S4** and **Figs. S6-S7**)^50,51^. The average S/S_0_ values decreased from 0.81 to 0.53 for β-1,3-glucan, from 0.47 to 0.35 for α-1,3-glucan, and from 0.53 to 0.46 for chitin in the mutant relative to the wild type. In the wild-type conidia, β-1,3-glucan is the most hydrated component, bridging to the mobile phase and forming a water-rich matrix; however, in the mutant it becomes similarly dehydrated to chitin and α-glucans, likely due to its tighter association with these molecules. Together, these results indicate that the denser packing of polymers in the Δ*sglA* cell wall more effectively excludes water, leading to uniformly low hydration across all polymers.

The molecular dynamics of the wall polymers further support this structural consolidation. Compared with the wild type, all polymers in the mutant exhibit longer relaxation time constants in both ^13^C-T_1_ (**Figure 3D**) and ^1^H-T_1_ (**Figure 3E**) measurements, indicating slower relaxation and reduced molecular motion across both the nanosecond and microsecond timescales (**Fig. S8** and **Table S5**). Despite this global rigidification, the different polysaccharides display distinct dynamic behaviors intrinsically. In both samples, β-1,3-glucan shows relatively short ^13^C-T_1_ values but longer ^1^H-T_1_ values, consistent with rapid local motions that are nevertheless constrained at larger length scales. In contrast, α-1,3-glucan shows the opposite trend, with little flexibility at the local scale, likely due to its extensive attachment to other wall polymers, but greater motion at the microsecond timescale, reflecting slow, collective movements within the rigid network to which it is integrated.

### Deletion of *sglA* in *A. fumigatus* conidia reshapes neutrophil responses

To better understand how the unique structural features of the Δ*sglA* conidial cell wall revealed by solid-state NMR influence host-pathogen interactions, we exposed *A. fumigatus* conidia *in vitro* to HL60 cells differentiated into neutrophils. We assessed both the activation state of the human cells and their ability to interact with the fungus (**Fig. 6A-C**). A modest increase in neutrophil activation was observed upon exposure to the Δ*sglA* mutant compared with the WT strain (**Fig. 6A, B**). More pronounced differences were detected when examining cell-fungus interactions. HL60 cells exposed to the Δ*sglA* mutant exhibited reduced interaction with fungal conidia, characterized by fewer adherent events and a higher proportion of fungus-free HL60 cells compared with WT exposure (**Fig. 6C**). Impaired interaction with the Δ*sglA* mutant diverts HL60 cells from efficient phagocytosis toward NETosis. Consequently, HL60 cells exposed to the Δ*sglA* mutant exhibit markedly increased NET formation (**Fig. 6D**), reduced fungal killing capacity (**Fig. 6E**), and decreased cell viability (**Fig. 6F**). Thus, SG accumulation and cell wall remodeling in the Δ*sglA* mutant lead to an increased capacity to trigger immune activation in HL60 cells *in vitro*.

Next, we examined the impact of this altered interaction *in vivo*, using an immunocompetent mouse model of invasive aspergillosis (**Fig. 6G**), thereby avoiding converting mice into immunodeficient, to assess the effect on neutrophilic response *in vivo*. At 7 days post-infection, mice exposed to the Δ*sglA* mutant exhibited significantly higher lung fungal burden and increased neutrophil recruitment compared with WT-infected mice (**Fig. 6H-J**). Despite these differences, overall survival did not differ between WT- and Δ*sglA*-infected mice. When measuring cytokine production in the lungs of Δ*sglA*-infected mice, we found a significant decrease in cytokines usually produced upon Dectin-1 interaction, including IL-27, IL-23, IL-17A, and TNF-α, compared with WT-infected mice (**Figure 4K**). In contrast, IL-12 levels were significantly higher in Δ*sglA*-infected mice compared with WT-infected mice (**Figure 4K**), consistent with increased engagement of the mutant strain with Toll-like receptor 2 (TLR2).

**Figure 4.**
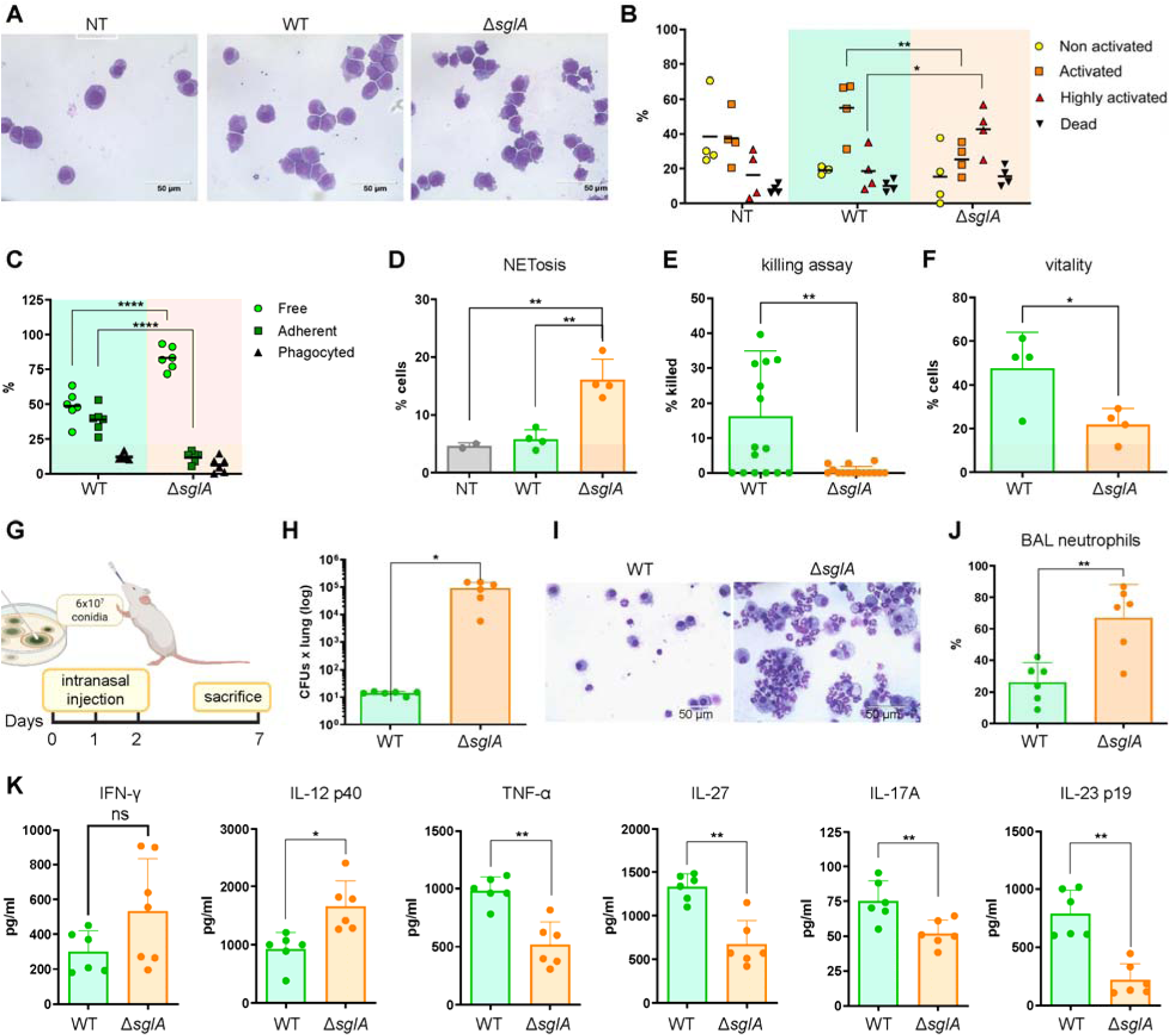
Effect of *A. fumigatus sglA* deletion on neutrophil response *in vitro* and *in vivo*. (**A**) Neutrophil-like differentiated HL60 cells either not-treated (NT) or stimulated with 1:1 ratio of *A. fumigatus* conidia (WT or Δ*sglA*) for 2h at 37°C. After stimulation, cells were cytospin and stained with May-Grünwald Giemsa. (**B**) Cell morphology and (**C**) interaction with *Aspergillus* conidia were evaluated. Non-activated cells were round; activated cells displayed an irregular shape and a reduced cytoplasm-to-nucleus ratio; highly activated cells exhibited blebbing and long cytoplasmic protrusions. Cells were considered dead when appearing small and anucleated. Conidia-cell interactions were quantified as the percentage of free (unbound), adherent (surface-associated), or internalized (phagocytosed) conidia. Statistical significance was determined by two-way ANOVA (*P<0.05, **P<0.01, ****P<0.0001). (**D**) Frequency of NETosis events. Statistical significance was determined by one-way ANOVA (**P<0.01). (**E**) Ability of neutrophil-like differentiated HL60 to eliminate *Aspergillus* conidia after 2h stimulation assessed with a killing assay. (**F**) Vitality of neutrophil-like differentiated HL60 after 24h stimulation with conidia at 1:1 ratio, expressed as percentage of the non-treated controls. (**G**) A murine model of invasive aspergillosis established through intranasal injection of 6 × 10^7^ WT or Δ*sglA Aspergillus* conidia for three consecutive days. (**H**) Lung CFUs counted after mice were sacrificed on day-7. (**I**) Bronchioalveolar lavage (BAL) recovered and fixed trough cytospin and stained with May-Grunwald Giemsa. (**J**) Neutrophil numbers in BAL counted and expressed as percentage of total cells. (**K**) Elisa test for IFN-γ, IL-12, TNF-α, IL-27, IL-17A and IL-23 performed on the lung homogenate of sacrificed mice. Statistical significance was determined by unpaired t-test (*P<0.05, **P<0.01).

## DISCUSSION

In this study, we show that deletion of *sglA* gene in *A. fumigatus* conidia triggers extensive remodeling of the cell wall, characterized by i) a marked increase in α-1,3-glucan within the rigid inner scaffold, ii) a compensatory decrease in β-1,3-glucan in this compartment, iii) complete loss of chitosan, iv) enhanced intermolecular contacts between α-1,3- and β-1,3-glucans, v) restricted molecular motions, and vi) reduced water accessibility. Together, these features may help preserve cell wall strength and integrity in the absence of *sglA*-dependent homeostasis. This tightened conidial cell-wall architecture plausibly contributes to the altered phenotype, in which Δ*sglA* conidia exhibit reduced germination capacity, shorter germ tubes, and delayed hyphal growth^30^. These structural features can be understood in the broader context of fungal cell wall organization and remodeling.

Notably, the reduction in chitin observed here in intact resting conidia is consistent with recent chemical analyses of Δ*sglA* cell-wall carbohydrates^30^. In that earlier study, β-glucan levels appeared unchanged relative to the wildtype^30^; however, our present data refine this interpretation by showing that β-glucan is undergoing a phase redistribution, with a fraction shifting from the rigid scaffold to the more mobile matrix. It is also important to note that the previous measurements were performed on mixed morphotypes, including germinating conidia, hyphae, and conidia that remained ungerminated^30^, whereas the current work focuses exclusively on resting conidia. Taken together, these complementary datasets suggest that β-glucan remodeling is stage-dependent, varying across distinct phases of fungal differentiation.

Recently, solid-state NMR and functional-genomics studies of *A. fumigatus* hyphae have defined a general architectural framework for the fungal cell wall, in which a rigid scaffold of chitin, β-1,3-glucan and α-1,3-glucan is embedded within a more mobile matrix enriched in galactomannan, glucans, and other biopolymers, such as galactosaminogalactan^48,52,53^. This bimodal organization is not static: under environmental and pharmacological stress, the wall undergoes coordinated remodeling in which the relative abundance and interactions of these polymers are dynamically rebalanced to preserve integrity^35,54,55^. These observations establish a key conceptual principle: the cell wall functions as a reconfigurable polysaccharide composite whose rigid core can be reinforced or redistributed in response to perturbation.

Building on this structural framework, conidial cell walls represent a developmentally specialized form of this architectural system. Solid-state NMR snapshots across morphotype transitions show that conidia possess a rigid, relatively dehydrated inner scaffold that undergoes defined polymer reorganization at the onset of germination^38^. Dormant conidia have a rigid core in which β-1,3-glucan contributes more strongly than α-1,3-glucan and chitin, whereas during swelling the amount of α-1,3-glucan temporarily doubles before returning toward its original proportion as germination progresses^38^. Developmental transitions are accompanied by increased surface accessibility of wall polysaccharides, including exposure of α-1,3-glucan during swelling and the appearance of mobile galactosaminogalactan at the surface of emerging germ tubes, thereby linking these early remodeling events to the structural principles that support the formation of mature hyphae at later developmental stages^38^. Consistent with this developmental specialization, Kre6-dependent β-1,6-glucan biosynthesis appears to be restricted to the conidial stage, reinforcing the view that polysaccharide composition and wall architecture are developmentally programmed and re-shaped during the transition to germination^56–58^.

Across these contexts, α-1,3-glucan is emerging as a central adhesive and buffering polymer. Although early studies suggested that α-1,3-glucan was dispensable, as synthase deletions caused only subtle growth or virulence phenotypes^59^, solid-state NMR later revealed that α-1,3-glucan is a major structural component that packs with chitin and β-1,3-glucan to form the stiff inner cell-wall core in *A. fumigatus*^48,52^. This supports a model in which α-1,3-glucan physically stabilizes interactions among wall polysaccharides and buffers architectural changes during morphogenesis and stress-induced remodeling^35,38,60^. Its adhesive role also provides a mechanistic explanation for earlier observations that surface-exposed α-1,3-glucan mediates aggregation of swelling conidia, an effect that will be abolished by treatment with α-1,3-glucanase^61,62^. The structural role of α-1,3-glucan in cell-wall construction is also evident in other fungi, such as *C. neoformans*, where it makes up the bulk of the rigid cell-wall core and interacts with essentially all other polysaccharides, along with melanin and the capsule^40^.

Previous *in vivo* studies demonstrated that Δ*sglA* conidia are efficiently cleared from immunosuppressed hosts while conferring complete protection against subsequent lethal wild-type challenge, in both live and heat-killed form^30^. In this study, the *in vitro* observations of reduced neutrophil-fungus interaction, impaired phagocytosis, diversion toward NETosis, and decreased fungal killing, along with the *in vivo* evidence of increased fungal burden, heightened neutrophil recruitment, and altered cytokine production, are consistent with a model in which cell wall remodeling in the Δ*sglA* mutant alters pathogen-associated molecular pattern (PAMP) exposure and disrupts normal host immune recognition.

Consistent with this interpretation, and at the molecular level, solid-state NMR analysis confirmed that the Δ*sglA* mutant exhibits a substantial increase in α-1,3-glucan content, including magnetically distinct α-1,3-glucan polymorphs that show extensive interactions with β-glucans. This expanded and well-integrated α-1,3-glucan fraction likely restricts and masks the associated β-1,3-glucan, reducing its accessibility to Dectin-1 during swelling, a stage at which β-1,3-glucan normally becomes exposed and recognized by innate immune cells, and thereby weakening β-glucan-dependent antifungal responses while redirecting immune signaling toward alternative pathways such as TLR2^63^. As reported for other opportunistic fungal pathogens, the presentation of α-1,3-glucan can effectively conceal β-1,3-glucan signatures and disrupt normal host immune recognition^64–66^. In this way, structural reorganization of the Δ*sglA* cell wall provides a mechanistic explanation for the defective immune recognition and altered neutrophil responses observed in both cellular and animal models.

Although the β-glucan content decreased in the rigid core, it remains plausible that the reduced Dectin-1 activity we observed occurred even though the β-glucan content increased in the mobile phase. This suggests that i) the mobile phase detected here does not necessarily correspond to the outer cell-wall layer, but instead reflects portions of the mobile matrix that neither aggregate nor associate with chitin; and ii) the mobile β-1,3-glucan is not responsible for Dectin-1 binding. Rather, only a subset of the more rigid β-1,3-glucan adopts the appropriate conformation for Dectin-1 recognition, consistent with reports that pattern-recognition receptors such as Dectin-1 bind most effectively to the triple-helical structure of β-glucan^67–69^. Together, these observations provide a plausible explanation for the reduced host immune response. Future work should correlate NMR-observed structural polymorphism with the diverse functional and immunological roles of cell-wall carbohydrates.

Taken together, our results support a model in which α-1,3-glucan acts both as an architectural adhesive and as a compensatory structural buffer in the Δ*sglA* mutant, stabilizing the rigid scaffold while retaining a substantial fraction of β-1,3-glucan through intermolecular interactions and thereby limiting its accessibility to host receptors during swelling. This organization provides a structural basis for the immunoprotective properties of the Δ*sglA* strain and parallels observations in *C. neoformans*, where related mutants show altered engagement of pattern-recognition receptors^30,70^. Future studies dissecting receptor-specific signaling will be important to define how these architectural changes shape the immune-protective profile of Δ*sglA* and related vaccine candidates.

## MATERIALS AND METHOD

### Preparation of uniformly ^13^C, ^15^N-labeled *A. fumigatus* cells for NMR analysis

Two *A. fumigatus* strains were used: the parental strain Δ*akuB*^KU80^, which enhances homologous recombination^42^, and the Δ*sglA* mutant, which has shown potential as a vaccine candidate. The strains were cultured on agar plates containing 20 g/L ^13^C-glucose (Catalog # CLM-1396-PK, Cambridge Isotope Laboratories) and a sodium nitrate salt solution (NLM-712-PK, Cambridge Isotope Laboratories) as the sole carbon and nitrogen sources, respectively, and supplemented with trace elements (**Table S6**). Cultures were incubated at 37 °C for 3 days. Conidia were collected from the plates using an aqueous 0.5% Tween-20 solution, washed twice with deionized water, followed by a wash with phosphate-buffered saline (PBS) to remove excess salts and glucose, and then centrifuged at 3000 rpm for 10 min. The intact conidia were packed into a 3.2-mm rotor or a 1.3 mm rotor for solid-state NMR analysis.

### SEM imaging of cell morphology

Fungal cultures were harvested and fixed in 4% (v/v) glutaraldehyde prepared in 0.1 M sodium phosphate buffer (pH 7.4) for 1-2 h at 4°C. Following primary fixation, samples were rinsed three times with phosphate-buffered saline (PBS) and post-fixed in 1% (w/v) osmium tetroxide for 1-2 h at room temperature. Specimens were then dehydrated through a graded ethanol series (25%, 50%, 75%, and 95%), with each dehydration step carried out for 10-15 min. Dehydrated samples were subjected to critical point drying using a Leica EM CPD300 with CO_2_ as the transitional fluid. The dried material was mounted onto aluminum stubs using conductive carbon tape. Scanning electron microscopy (SEM) was performed using a JEOL JSM-7500F field-emission instrument, and micrographs were obtained at multiple magnifications to evaluate fungal surface morphology.

### ^13^C solid-state NMR analysis of polysaccharides present in conidial cell wall

High-resolution solid-state NMR spectroscopy was performed on a Bruker Avance Neo 800 MHz spectrometer equipped with a 3.2 mm HCN triple-resonance MAS probe at the Max T. Rogers NMR Facility, Michigan State University. ^13^C-detected experiments were carried out at a magic-angle spinning (MAS) frequency of 15 kHz and a regulated sample temperature of 298 K. Chemical shifts were externally referenced to the tetramethylsilane (TMS) scale by calibrating the methylene (CH_2_) resonance of adamantane to 38.48 ppm. Radiofrequency (rf) field strengths for ^1^H hard pulses, heteronuclear decoupling, and cross-polarization (CP) transfers ranged from 71.4 to 83.3 kHz. ^13^C pulses were applied using radiofrequency field strengths of 50 or 62.5 kHz, depending on the specific experiment.

Resonance assignments of carbon sites within polysaccharides were obtained using a series of 2D solid-state NMR experiments designed to probe the molecular organization of the fungal cell wall. Through-bond ^13^C-^13^C connectivities of mobile components were characterized using ^13^C-DP refocused J-INADEQUATE experiments carried out with a short 2-s recycle delay^44,71^. The J-evolution period consisted of four delays of 2.3 ms each, optimized for achieving the highest carbohydrate intensity. In parallel, rigid polysaccharides were analyzed using CP-based 2D correlation experiment using a 53 ms CORD (COmbined R2nv-Driven) mixing period at 13.5 kHz MAS (**Fig. S2**)^43,72^. The ^13^C signals of mobile and rigid carbohydrates were assigned based on carbon connectivities in the J-INADEQUATE spectra and intramolecular cross-peaks in the CORD spectra, and were cross-validated using data from the Complex Carbohydrate Magnetic Resonance Database and recent literature reports^73^. The chemicals shifts are documented in **Table S7.**

### Compositional analysis of cell wall carbohydrates

Analysis of peak volumes from 2D ^13^C-CP CORD and 2D ^13^C-DP refocused J-INADEQUATE spectra allowed us to estimate the molar composition of rigid and mobile carbohydrates in each sample, respectively (**Tables S2**-**3**). Peak volumes were integrated using the Bruker TopSpin software package (version 4.1.4). To minimize errors arising from spectral overlap, only well-resolved and unambiguous resonances were included in the quantitative analysis. For the CORD spectra, analysis was performed by averaging the volumes of clearly resolved cross-peaks associated with each polysaccharide component. For the INADEQUATE spectra, only well-defined scalar-coupled carbon pairs were considered. Relative molar abundances were calculated by normalizing the integrated peak volumes to the number of contributing resonances for each carbohydrate species. Standard errors were estimated by dividing the standard deviation of the integrated volumes by the number of cross-peaks included in the analysis. Total standard error for each sample was obtained by calculating the square root of the sum of the squared individual errors^36^.

### Profiling the hydration and mobility of cell wall polysaccharides

The dynamics of cell wall polysaccharides were analyzed using two relaxation methods. The ^13^C-T_1_ relaxation times were measured using the Torchia-CP pulse sequence with z-filter durations ranging from 0.1 µs to 12 s. For each resolved resonance, the decay in signal intensity was monitored as the z-filter duration increased, and the resulting curves were fit to a single-exponential function to obtain the ^13^C-T_1_ relaxation time constants. Absolute intensities were pre-normalized to the number of scans collected for each spectrum. Similarly, ^13^C-detected ^1^H-T_1ρ_ relaxation times were measured using a Lee-Goldburg (LG) spinlock with varying duration combined with LG-CP. This approach effectively suppressed ^1^H-^1^H spin diffusion during both the spinlock and CP periods, enabling site-specific determination of ^1^H-T_1ρ_ relaxation by detecting the directly bonded ^13^C sites^74^. Peak intensity decays were modeled using a single-exponential function to extract the ^1^H-T_1ρ_ time constants. All relaxation datasets were processed and analyzed using Origin 2021.

To probe polysaccharide hydration levels, water-edited 2D ^13^C-^13^C correlation spectra were acquired^50,51,75,76^. The experiment began with ^1^H excitation, followed by a ^1^H-T_2_ filter (0.45 ms × 2 for the WT sample and 0.6 ms × 2 for the mutant), which eliminated 97% of polysaccharide proton signals while retaining 84% of bulk water magnetization. Water-derived magnetization was then transferred to polysaccharides using a 4 ms ^1^H mixing step, followed by a 1-ms ^1^H-^13^C cross-polarization (CP) period for site-specific ^13^C detection. A 50 ms DARR mixing period was applied to both the water-edited spectrum and a corresponding control 2D spectrum that preserved full signal intensity. Hydration levels were quantified by calculating the relative intensity ratios between the water-edited (S) and control (S_0_) spectra for all cell wall samples. Signal intensities were normalized to the number of scans collected for each dataset before analysis. The key experimental parameters are summarized in **Table S8**.

### ^1^H-detected solid-state NMR resolving polymorphism and intermolecular interactions

Rigid molecules in *A. fumigatus* cell walls were characterized using CP-based ^1^H-detected experiments on a Bruker Avance Neo 600 MHz spectrometer located at the Max T. Rogers NMR Facility at Michigan State University with a fast-MAS 1.3 mm HCN triple-resonance probe spinning at 60 kHz. ^13^C chemical shifts were externally referenced to the TMS scale, and ^1^H chemical shifts were referenced to the DSS scale. Rigid polysaccharide regions were investigated using two ^1^H-detected experiments, including 2D hCH and 2D hChH with RFDR-XY16 mixing^47,77,78^. One-bond ^13^C-^1^H correlations were obtained using the 2D hCH experiment via a short second CP contact time of 50 µs, while through-space correlations were generated using the 2D hChH experiment via a ^1^H-^1^H RFDR-XY16 homonuclear dipolar recoupling period with a mixing time of 0.533 ms. The 90° pulse lengths were 2.5 µs (100 kHz) for ^1^H and 4 µs (62.5 kHz) for ^13^C. slpTPPM decoupling was applied on the ^1^H channel during t evolution^79^, with a radiofrequency field strength of 12.8 kHz, and WALTZ-16 decoupling was applied on ^13^C during proton detection at 20.2 kHz. Water suppression was achieved using the MISISSIPPI sequence (16 kHz, 100 ms). For both experiments, 448 TD points were acquired with 32 scans per increment and a recycle delay of 2 s, resulting in total acquisition times of 8 h 34 min (hCH) and 8 h 28 min (hChH). All multidimensional datasets were collected using the States-TPPI method^80^. Detailed experimental parameters and assigned ^13^C and ^1^H chemical shifts are provided in **Tables S9-S10**.

Mobile regions of the *A. fumigatus* conidial cell walls were investigated using J-coupling-based proton-detected experiments on a Bruker Avance Neo 800 MHz spectrometer equipped with a 3.2 mm HCN triple-resonance MAS probe operating at 15 kHz. The mobile molecules do not require fast MAS, as their intrinsic dynamics average out a significant portion of the ^1^H-^1^H dipolar coupling^81^. Mobile polysaccharides were assigned using a 2D ^13^C-^1^H correlation experiment J-hCcH TOCSY (total correlation spectroscopy) and 3D ^13^C-^13^C-^1^H correlation experiment J-hCCH TOCSY employing DIPSI-3 (decoupling in the presence of scalar interactions) mixing^49,82^. The 90° pulse widths were set to 3.5 µs (71.4 kHz) for ^1^H and 5.0 µs (50 kHz) for ^13^C. SPINAL-64 (small phase incremental alternation with 64 steps) heteronuclear decoupling was applied during the t_1_ and t_2_ evolution periods with an rf field strength of 71.4 kHz, while WALTZ-16 (wideband alternating-phase low-power technique for zero-residual splitting) decoupling was applied on ^13^C during ^1^H detection with an radiofrequency field strength of 17 kHz^83,84^. Water suppression was achieved using the MISSISSIPPI (multiple intense solvent suppression intended for sensitive spectroscopic investigation of protonated proteins) sequence with a 26 kHz radiofrequency field applied for 40 ms^85^. Broadband DIPSI-3 mixing was employed to obtain ^13^C-^13^C correlations using a 2 ms spin-lock pulse. The mixing time was set to 25.5 ms, with a radiofrequency field strength of 17 kHz applied for both the DIPSI-3 and spin-lock pulses.

For the wild-type *A. fumigatus* conidial sample, the 3D hCCH TOCSY experiment was acquired with 128 time-domain (TD) points in both the t_1_ and t_2_ dimensions. Eight transients were co-added per TD point using a recycle delay of 2 s, resulting in a total experimental time of 3 days, 5 h, and 53 min. The 2D hcCH TOCSY experiment with DIPSI-3 mixing was performed on both samples using the same parameters as the 3D experiment, except t_1_ was set to a single TD point and t_2_ to 512 TD points. Thirty-two scans were acquired per increment with a recycle delay of 2 s, yielding a total experiment time of 9.6 h (**Tables S11**-**S12**).

### HL60 culture and differentiation

After thawing, HL60 were cultured into tissue-treated flasks maintaining a concentration between 1 × 10^5^ and 5 x 10^5^ cell/mL splitting every 2 to 3 days. The culture medium consisted of RPMI supplemented with 10% fetal bovine serum (FBS), 1% penicillin-streptomycin, and 1% GlutaMAX (Invitrogen). For differentiation into neutrophil-like HL60 cells, cells were stimulated by adding 1.25% DMSO to the culture medium for 7 to 10 days^86^. To check differentiation status, cytofluorimetry was performed to measure increased expression of surface marker CD16b compared to non-treated HL60.

### HL60 and *Aspergillus* interaction assay

Neutrophil-like HL60 cells (5 × 10^5^) were incubated in 1 mL of culture medium either without stimulus or with *A. fumigatus* conidia at a 1:1 cell-to-conidium ratio for 2 h at 37°C 5% CO_2_. Following incubation, 50 μL of cell suspension was recovered, and cells were cytospinned into microscopy slides through a cytospin chamber at 700 × g for 7 min and minimum rotor acceleration and deceleration. Produced slides were stained with May-Grunwald-Giemsa staining (Sigma-Aldrich). Cell morphology, interaction between cells and conidia and netosis events were measured, counting 10 fields at 60-fold magnification for each slide on an EVOS® FL Auto Imaging System (Thermo-Fisher Scientific). Cells undergoing NETosis were distinguished by the presence of densely Giemsa-stained smears.

### HL60 vitality assay

Neutrophil-like HL60 cells (5 × 10^5^) were incubated in 1 mL of culture medium either without stimulus or with *A. fumigatus* conidia at a ratio of 1:1 for 24 hours at 37°C 5% CO_2_. After the incubation, vitality was assessed using trypan blue solution and a haemocytometer.

### *Aspergillus* killing assay

Conidia of *A. fumigatus* (5 × 10^5^), both wild-type and Δ*sglA*, were incubated in 100 μL of culture medium for 2 h at 37°C 5% CO_2_ with and without neutrophil-like HL60 cells at a ratio of 1:1. Following incubation, 10 μL of TRITON 100X was added to each well and mixed vigorously. The plate was left to incubate for 15 min at 37°C to lyse the HL60 cells. Each well was diluted 1 to 5000 into PBS/tween20 0,05%. 100 μl of each solution was seeded in Sabouraud agar plates (Merk Millipore) and left to incubate overnight at 37°C. *A. fumigatus* CFUs were counted, and the killing percentage was expressed as the difference between conidia not exposed to phagocytic cells and those exposed.

### Murine model of invasive aspergillosis

Experimental protocols for murine in vivo studies were approved by the Ministry of Health (Authorization N. 310/2025-PR) and previously certified by the animal ethics committee ‘OPBA’ from the University of Perugia, Italy. All mice used in this study were female C57BL/6 mice, 8-10 weeks old, purchased from Charles River Mice were anesthetized by intraperitoneal (i.p.) injection of 2.5% Avertin (Sigma Chemical Co.) before intranasal instillation of 6×10^7^ *A. fumigatus* resting conidia suspended in 20 µL of saline, administered once daily for three consecutive days. Mice were euthanized on day 7. Bronchioalveolar lavage (BAL) was performed on sacrificed animals by cannulating the trachea and washing the airways with PBS to collect the BAL fluid. Differential cell counts were generated on BAL smears stained with May-Grünwald Giemsa (Sigma-Aldrich) reagents, counting 10 fields at 60X magnification on an EVOS® FL Auto Imaging System (Thermo-Fisher Scientific). Lungs recovered from sacrificed mice and homogenized in 1 ml of PBS. 100 μl of homogenate was seeded in Sabouraud agar plates (Sigma-Aldrich) and incubated at 37°C overnight for CFU counting. Homogenate was centrifuged at 2000 rpm for 10 min, and supernatant was recovered for ELISA test. ELISA tests for IFN-γ (Biolegend), IL-12 p40 (eBioscience), TNF-α, IL-27, IL-17A, IL-23 p19 (Invitrogen) were performed as per the producers’ instructions.

## DATA AVAILABILITY

All relevant data that support the findings of this study are provided in the article and Supplementary Information. All the original ssNMR data files will be deposited in the Zenodo repository and the access code and DOI will be provided.

## AUTHOR CONTRIBUTIONS

K.S., and A.A. conducted ^13^C solid-state NMR experiments. A.A. and J.R.Y. conducted ^1^H-detection solid-state NMR experiments. K.S., A.A., and T.W. analyzed solid-state NMR data. K.S and C.M.F. prepared fungal samples. G.V., A.S. and T.Z. performed experiments *in vitro* with neutrophils and *in vivo* expreimentation. A.A. and K.S. wrote the first draft. All authors wrote the manuscript. M.D.P. and T.W. designed and supervised the project.

## ACKNOWLEDGMENT

The research was supported by the National Institute of Health (NIH) under award numbers R01AI173270 to T.W. and R01AI125770 to M.D.P.

