## Supplementary file for "α-1,3-Glucan-Driven Remodeling of the Conidial Cell Wall in an *Aspergillus fumigatus* Vaccine Strain Alters Innate Immune Recognition"

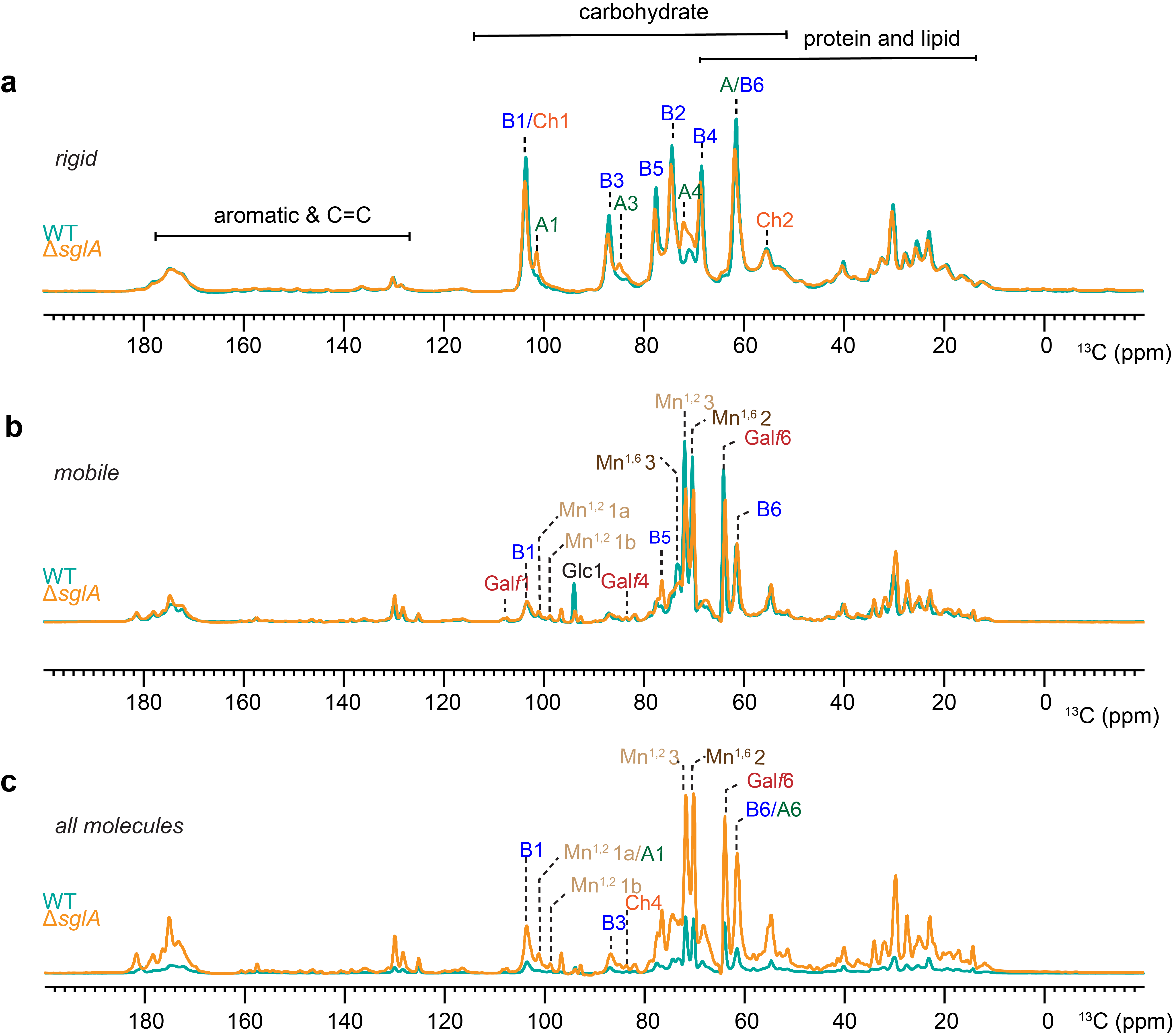

**Figure S1.** **1D ^13^C spectra of intact *A. fumigatus* conidia.** The spectra are color coded to show the cyan spectra for WT and yellow for mutant Δ*sglA*. (**a**) Carbohydrates showing the rigid glucans obtained by CP experiment. (**b**) The mobile glucans probed by 2 s DP spectra and (**c**) The quantitative glucans represented by the 35 s DP spectra.

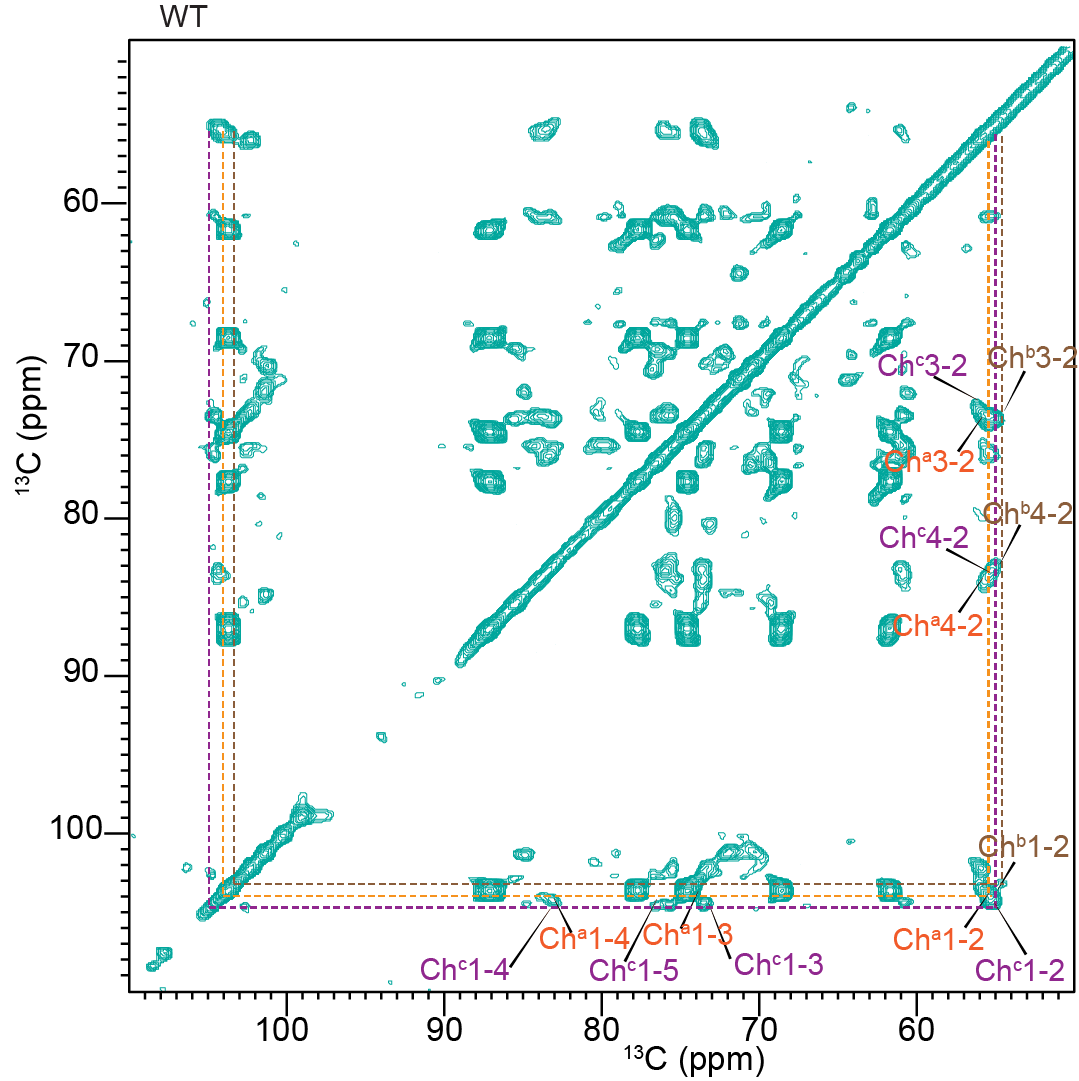

**Figure S2.** **2D ^13^C-^13^C CORD spectra showing the rigid components in *A. fumigatus* WT conidia.** It shows the presence of polymorphic forms of chitin.

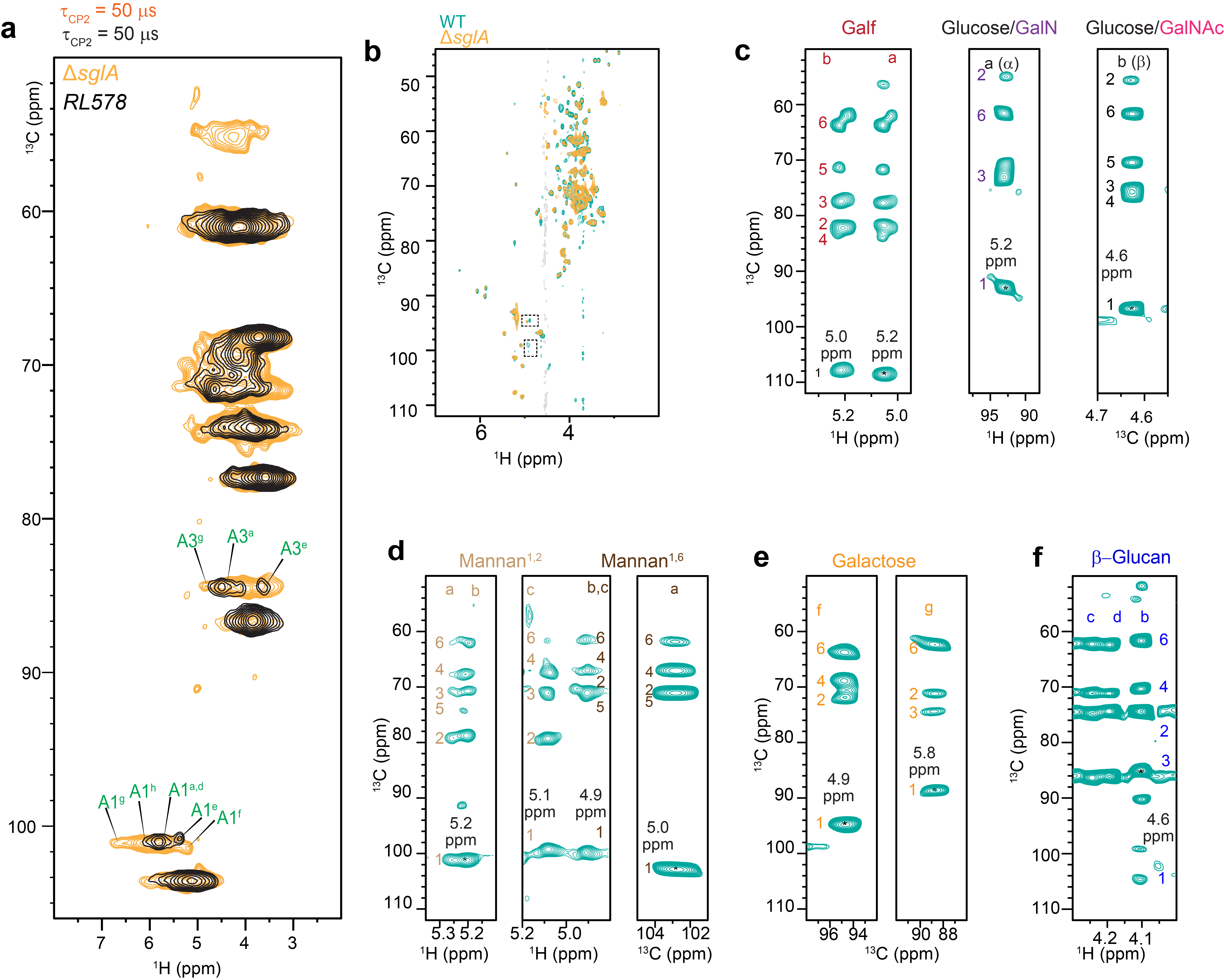

**Figure S3.** **Identifying polymorphic forms of rigid polysaccharides in of *A. fumigatus* conidial wall.** (**a**) Comparison of the resolved polymorphic forms of α-1,3-glucan in the fully protonated conidial cell wall of mutant Δ*sglA* and the partially deuterated mycelial cell wall of the *RL578* wild-type cells. The polymorphic forms a (^13^C:101.1 and ^1^H:5.7 ppm), d (^13^C: 101.1 ppm, ^1^H: 5.7 ppm), e (^13^C: 100.76 ppm, ^1^H: 5.4 ppm) and h (^13^C: 101.1 ppm, ^1^H: 6.0 ppm) were matched. Additionally, two novel forms, designated as f (^13^C: 101.42 ppm, ^1^H: 5.2 ppm) and g (^13^C: 101.0 ppm, ^1^H: 6.58 ppm) were detected in ∆*sglA* mutant. (**b**) Overlaid carbohydrate regions of the 2D hcCH TOCSY (DIPSI-3) spectra of *A. fumigatus* wild-type (WT) (green) and the ∆*sglA* (mutant) (orange) cell walls, showing one-bond ^13^C-^1^H correlations. Water singles are shown in grey. Dotted squares highlight signals present in the WT but absent in the ∆*sglA* mutant, indicating specific carbohydrate components affected by the gene deletion. The strips were extracted from the 3D hCCH TOCSY (DIPSI-3) spectrum and show through-bond carbon-carbon connectivity. The strip extracted at a specific carbon or proton site is indicated by an asterisk (*). Resolved signals assigned to: (**c**) two forms of glucofuranose (Gal*f*), one form of galactosamine (GalN), and one form of N-acetyl galactosamine (GalNAc); (**d**) three forms each of α-1,2-mannan and α-1,6-mannan (**e**) seven forms of galactose derivatives; (**f**) four forms of β-glucan; Carbon positions are numbered (1-6), and lowercase letters denote different allomorphic forms. All spectra of were acquired on a 14.1 T spectrometer at a MAS rate of 60 kHz and (c) were on 800MHz at MAS of 15 kHz.

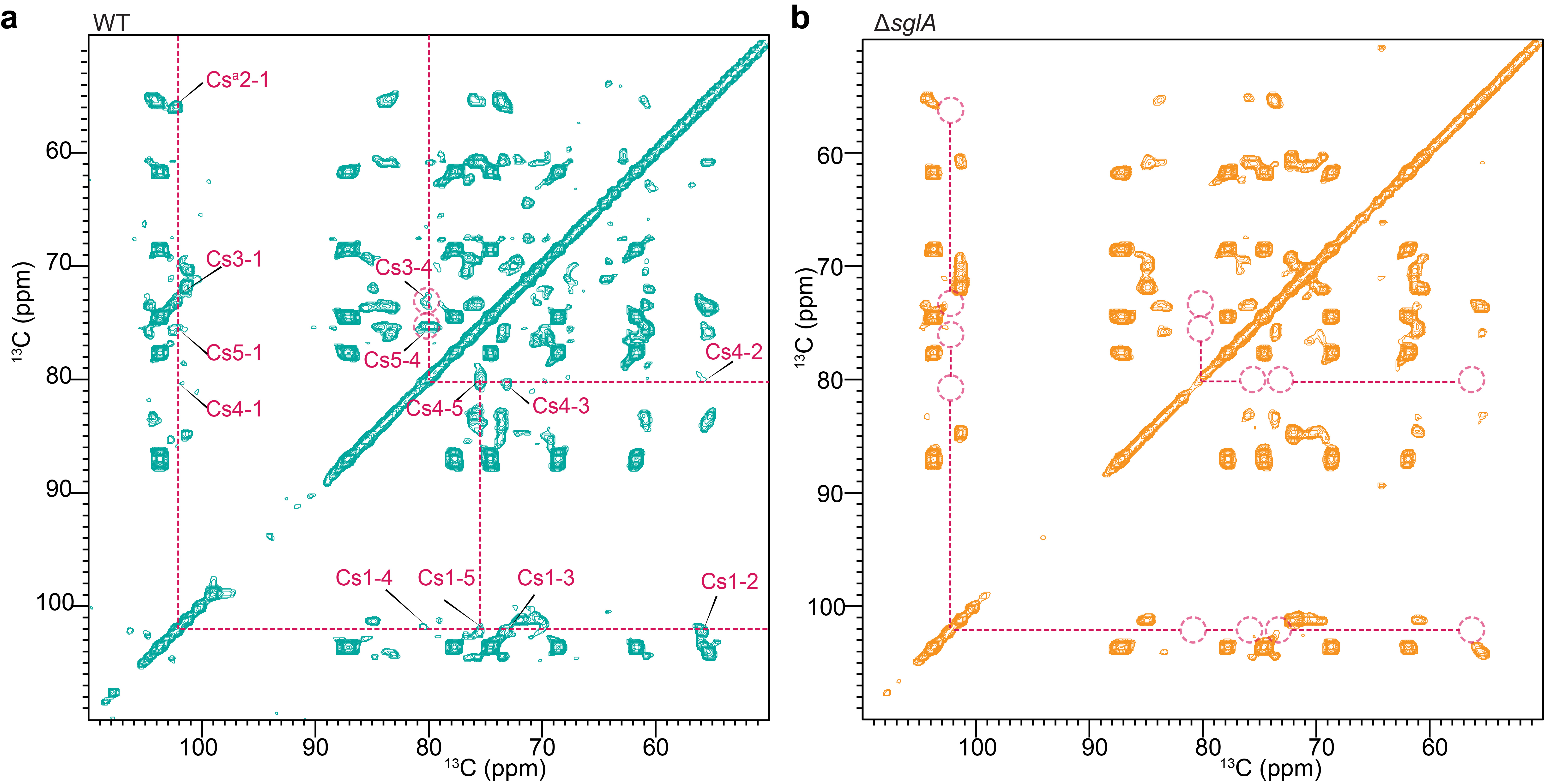

**Figure S4.** (**a**) 2D ^13^C-^13^C CORD spectra showing the rigid components in *A. fumigatus* WT conidia (cyan). It shows the presence of chitosan. (**b**) 2D ^13^C-^13^C CORD spectra showing the rigid components for mutant Δ*sglA* in orange spectra. It shows the absence of chitosan. All measurements were obtained using 800 MHz NMR spectrometer at 15 kHz.

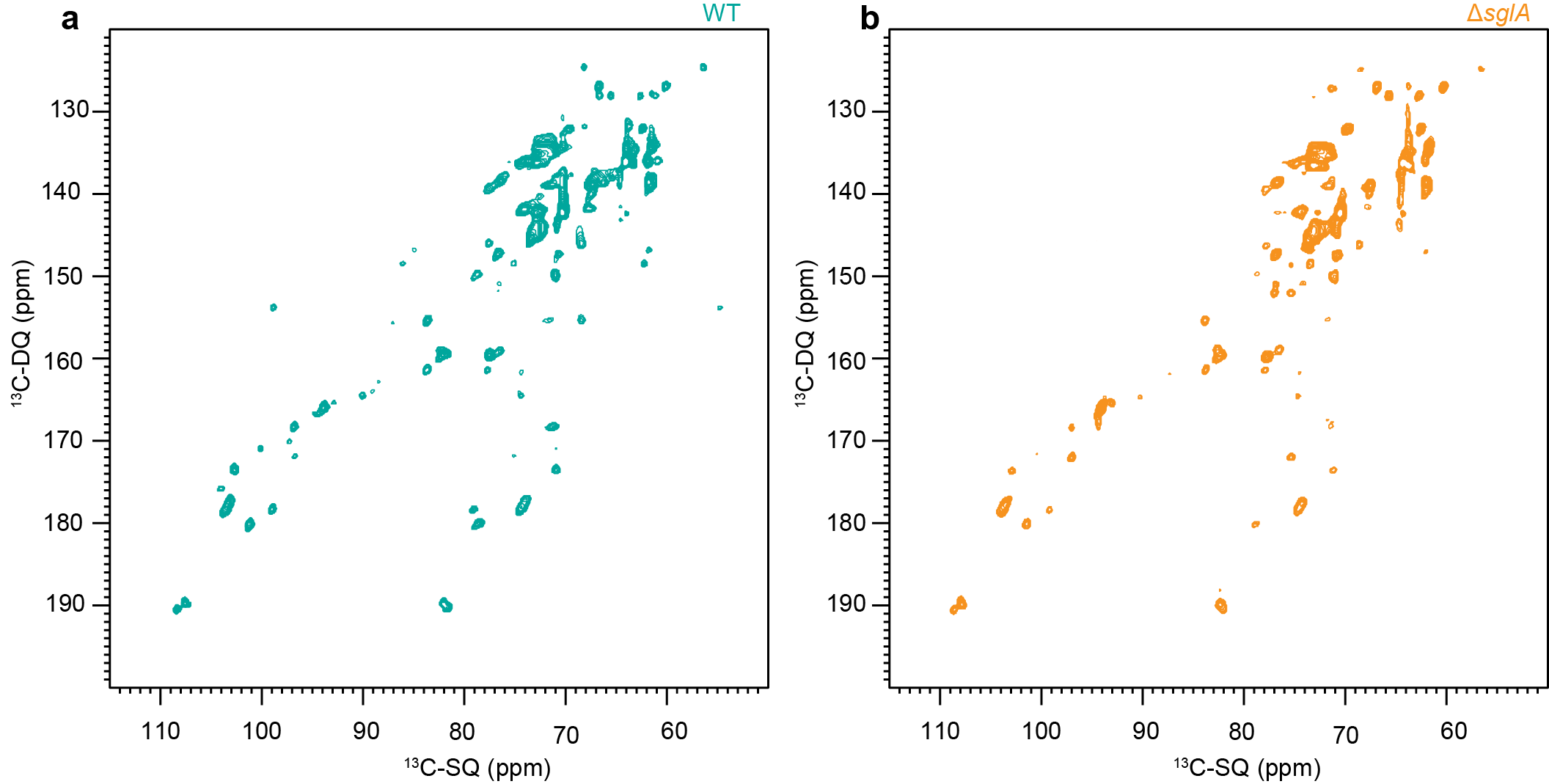

**Figure S5.** **Mobile carbohydrates in *Aspergillus* conidia.** (**a**) The 2D ^13^C-DP INADEQUATE spectra of WT conidia showing all the mobile glucans. (**b**) The mobile spectra representing mutant Δ*sglA* showing decrease in the intensity of mannose 1,2. The spectra of WT and mutant Δ*sglA* were measured on 800MHz at 15 kHz.

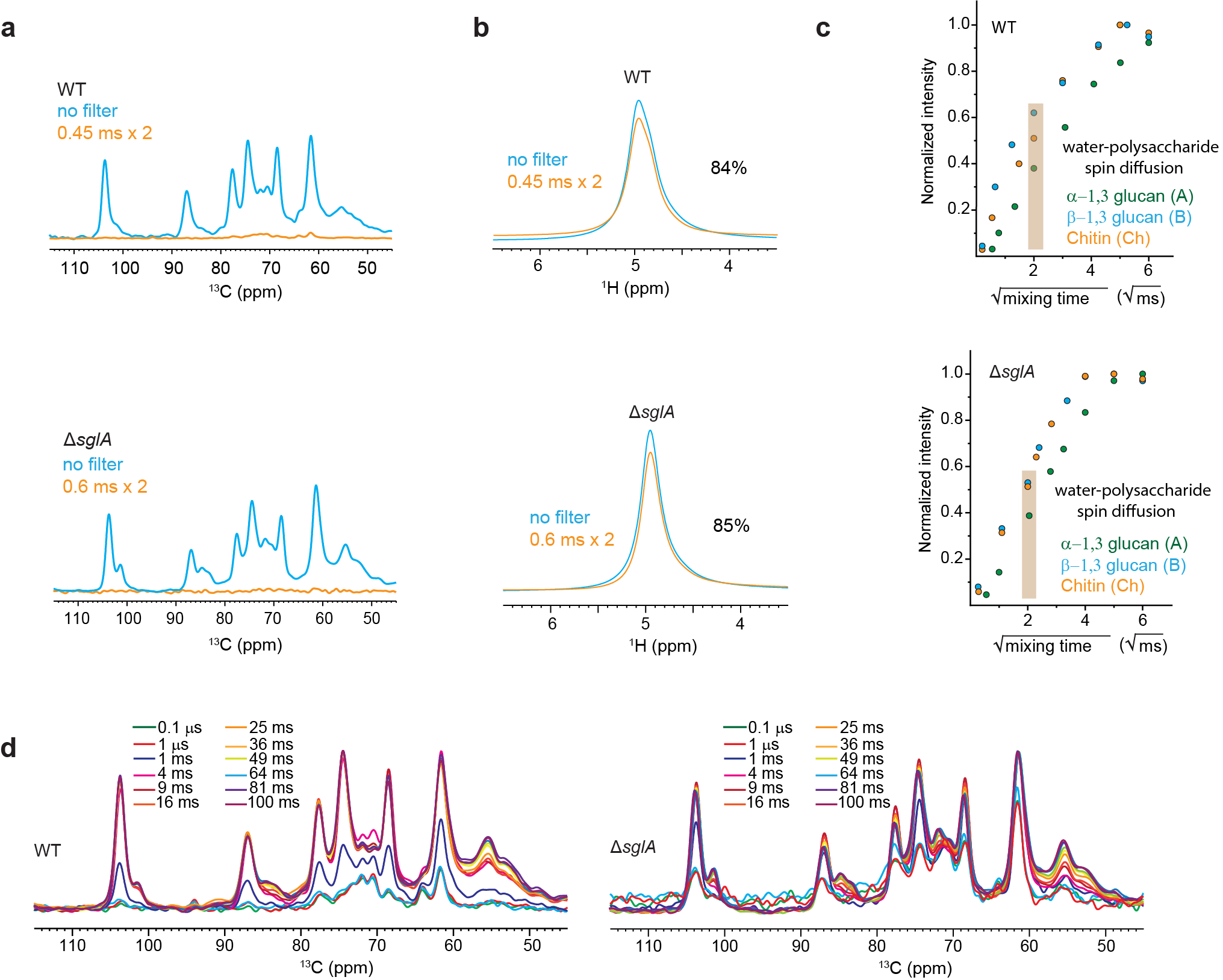

**Figure S6. Water-edited experiments to monitor carbohydrate-associated water. (a)** ^1^H-T_2_ filtered (orange) and control(blue) ^13^C spectra are shown for WT (top) and mutant (bottom). **(b)** ^1^H-T_2_ filtered (orange) and control (blue) ^1^H NMR spectra, with 84% and 85% of water signal retained for both samples after the ^1^H-T_2_ filter**. (c)** Representative water-to-polysaccharide ^1^H spin diffusion buildup curves. **(d)** 1D water-edited spectra with different ^1^H mixing times are shown for WT and mutant samples. All the spectra were measured on 400MHz spectrometer at 10 kHz at 277K.

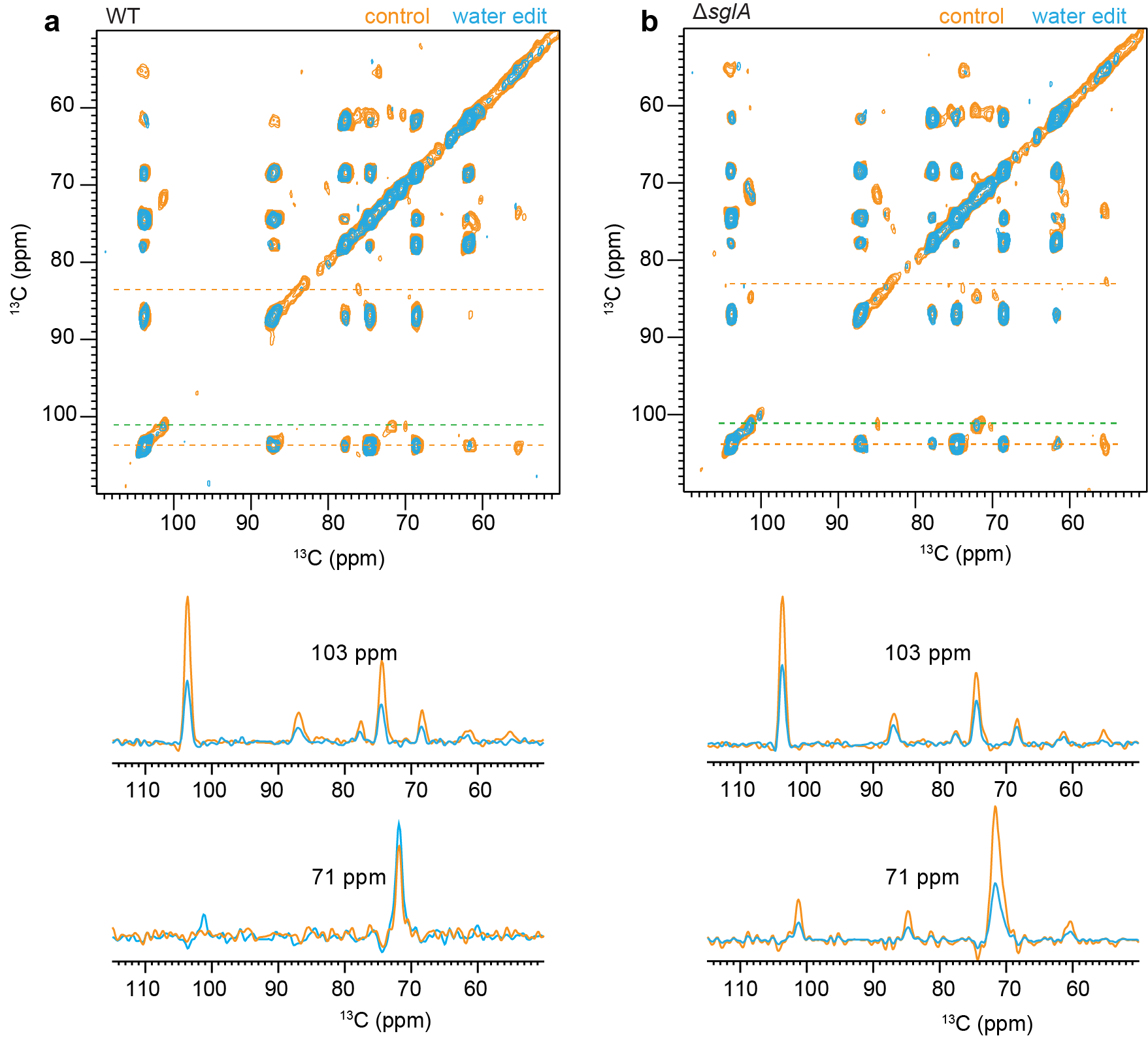

**Figure S7.** **Water-edited analysis for *Aspergillus* fumigatus conidial cell wall.** (**a**) Overlay of 2D water-edited (blue) and control (orange) ^13^C-^13^C correlation spectra (T2 = 0.45 ms × 2 for Sample. Representative 1D slices extracted from the 2D ^13^C-^13^C correlation spectra are shown for WT (left bottom) sample (**b**) Overlay of 2D water-edited (blue) and control (orange) ^13^C-^13^C correlation spectra (T_2_ = 0.6 ms × 2 for Sample. Representative 1D slices extracted from the 2D ^13^C-^13^C correlation spectra are shown for mutant (right bottom) sample All the measurements were taken on a 400 MHz spectrometer at 10 kHz MAS and 277 K.

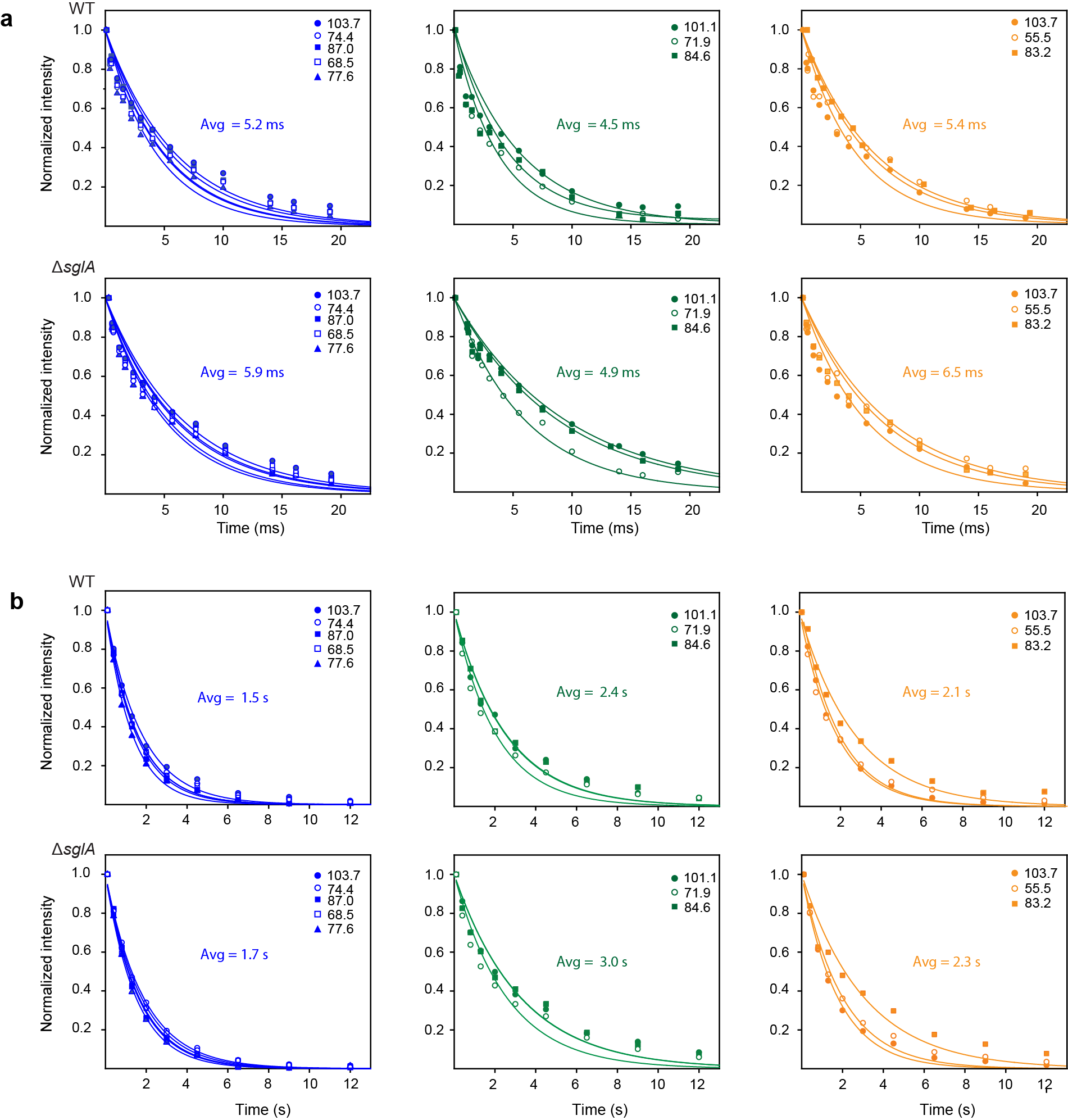

**Figure S8.** **Relaxation curves of polysaccharides in *A. fumigatus* conidial cell walls.** The relaxation decay curves are shown separately for (**a**) ^1^H-T_1ρ_ and (**b)**^13^C-T_1_ of the *A. fumigatus* WT and mutant Δ*sglA* conidia.

**Table S1.** **Average cell thickness of *A. fumigatus* conidial cells.** The results are the average and standard deviation of measurements from SEM images of different cells (n=9 for WT and Δ*sglA*, n=9).

| Strain | Average cell thickness (μm) |
| --- | --- |
| WT | 1.8 ± 0.2 |
| Δ*sglA* | 1.9 ± 0.2 |

**Table S2.** **Molar composition of rigid polysaccharides in *A. fumigatus* conidial cell wall.** The numbers were calculated using the integrals of well-resolved cross peaks of β-1,3 glucan, α-1,3 glucan, β-1,6 glucan, chitosan and chitin in 2D ^13^C-^13^C CORD spectra. The results were already normalized by the number of scans.

| Sample | α-1,3-glucan | β-1,3-glucan | Chitin | β-1,6-glucan | Chitosan |
| --- | --- | --- | --- | --- | --- |
| WT | 2 ± 1% | 78 ± 12% | 11 ± 3% | 4 ± 0.9% | 5 ± 1% |
| Δ*sglA* | 20 ± 4% | 67 ± 10% | 8 ± 2% | 5 ± 2% | NA |

The area of the following well-resolved cross peaks 53 ms CORD spectra are used:

β-1,3 (a): the average of C1-C2/3/4/5 and C3-C2/4/5/6.

β-1,6: the average of C3/5-C4 and C5-C6.

Chitin: the average of C1-2/4/5, C3-C2, C4-C2/3/5, and C5-C2.

Mannan: the average of C2-C5 and C2-C6.

**Table S3**. **Molar composition of mobile polysaccharides in *A. fumigatus* conidial cell wall.** The numbers were calculated using the integrals of well-resolved cross peaks of β-1,3 glucan, mannose units, and glucofuranose units, in ^13^C DP *J*-INADEQUATE spectra. The results were already normalized by the number of scans.

| Sample | β-1,3-glucan | Mn^1,2^ | Mn^1,6^ | Gal*f* |
| --- | --- | --- | --- | --- |
| WT | 22 ± 5% | 26 ± 4% | 22 ± 3% | 30 ± 8% |
| sglA | 46 ± 6% | 17 ± 2% | 13 ± 3% | 24 ± 8% |

The area of the following well-resolved cross peaks of 2D ^13^C *J*- INADEQUATE spectra are used:

β-1,3-glucan: the average of C1-C2, C2-C3 and C5-C6

Gal*f*: the average of C1-C2, C2-C3, C3-C4 and C4-C5

Mannan: the average of C1-C2, C2-C3 and C3-C4

**Table S4. Water-edited intensities of the polysaccharides**. The intensity ratios are obtained by comparing the peak intensities between water-edited and the control 2D spectra of the *Aspergillus* conidial cell walls. The error bars represent propagated standard deviations from NMR signal-to-noise ratios.

| β-1,3-glucan | | | | α-1,3-glucan | | | | Chitin | | | |
| --- | --- | --- | --- | --- | --- | --- | --- | --- | --- | --- | --- |
| (ω1 ppm, ω2 ppm) | | WT (S/S_0_) | Δ*sglA* (S/S_0_) | (ω1 ppm, ω2 ppm) | | WT (S/S_0_) | Δ*sglA* (S/S_0_) | (ω1 ppm, ω2 ppm) | | WT (S/S_0_) | Δ*sglA* (S/S_0_) |
| 103 ppm | 103.6  86.7  77.4  74.3  68.1  61.1 | 0.71±0.03  0.82±0.15  -  0.81±0.06  0.89±0.19  0.75±0.35 | 0.43±0.02  0.51±0.13  0.54±0.19  0.48±0.04  0.51±0.12  0.68±0.09 | 101 ppm | 101.6  84.6  71.6  69.6  60.2 | 0.35±0.08  0.23±0.09  0.44±0.07  0.64±0.07  0.85±0.11 | 0.38±0.08  0.34±0.05  0.35±0.07  -  - | 103 ppm | 103.6  55.3  73.1  76.3  82.5 | 0.71±0.07  0.31±0.01  0.29±0.01  0.5±0.0  0.36±0.02 | 0.43±0.02  0.32±0.05  -  -  0.61±0.14 |
| 86 ppm | 103.5  86.6  77.3  74.3  68.2  61.1 | 0.81±0.12  0.79±0.08  0.86±0.13  0.84±0.11  0.78±0.12  - | 0.62±0.10  0.47±0.06  0.67±0.12  0.52±0.08  0.51±0.09  0.86±0.10 | 84 ppm | 101.1  84.5  71.6  69.3  60.2 | 0.41±0.03  0.24±0.10  0.57±0.03  0.61±0.05  0.37±0.6 | 0.59±0.12  0.40±0.02  -  0.31±0.08  0.24±0.07 | 83 ppm | 103.6  55.3  73.6  75.9  83.4 | 0.13±0.03  0.35±0.02  0.50±0.05  0.56±0.05  0.19±0.03 | -  0.60±0.01  -  0.45±0.09  0.41±0.06 |
| 77 ppm | 103.5  86.71  77.1  74.4  68.1  61.2 | 0.94±0.4  0.74±0.26  0.98±0.08  0.88±0.47  0.79±0.10  - | 0.59±0.18  0.34±0.16  0.52±0.04  0.56±0.11  0.53±0.07  0.51±0.06 | 71.4 ppm | 101.2  84.7  71.6  60.3  69.3 | 0.42±0.18  0.19±0.02  0.48±0.05  0.58±0.06  0.63±0.09 | 0.16±0.05  0.34±0.02  0.65±0.006  0.43±0.08  0.35±0.06 | 55 ppm | 103. 3  54.7  73.6  75.4  83.4 | 0.59±0.03  0.31±0.01  0.42±0.01  0.68±0.09  0.74±0.02 | 0.55±0.14  0.32±0.06  -  -  0.42±0.11 |
| 74 ppm | 103.5  86.6  77.2  74.2  68.2  61.0 | 0.84±0.10  0.87±0.1  -  0.68±0.11  0.84±0.12  0.72±0.10 | 0.49±0.04  0.55±0.11  0.48±0.11  0.49±0.11  0.52±0.13  0.50±0.14 |  |  |  |  |  |  |  |  |
| 68 ppm | 103.6  86.6  77.4  74.3  68.5  61.1 | 0.74±0.15  0.82±0.17  0.77±0.13  0.60±0.11  0.74±0.04  0.74±0.14 | 0.51±0.12  0.49±0.11  0.53±0.07  0.52±0.10  0.49±0.03  0.43±0.11 |  |  |  |  |  |  |  |  |
|  | n | 26 | 30 |  |  | 15 | 12 |  |  | 15 | 9 |
| Average | | 0.81 | 0.53 |  |  | 0.47 | 0.35 |  |  | 0.53 | 0.46 |

**Table S5**. **^13^C-T_1_ and ^1^H-T_1ρ_ relaxation time constants of polysaccharides in *A. fumigatus.*** A single exponential equation was used to fit the T1 data 𝐼(𝑡) = 𝑒 ^−𝑡/𝑇^_𝐼_. A single exponential equation was used to fit the T_1ρ_ data: 𝐼(𝑡) = 𝑒 ^−𝑡/𝑇^_𝐼𝜌_. Error bars are standard deviations of the fit parameters.

| Sample type | Cross peaks | T_1_ (s) | Average | Cross peaks | T_1ρ_ (ms) | Average |
| --- | --- | --- | --- | --- | --- | --- |
|  | B1 | 1.8±0.1 |  | B1 | 6.1±0.5 |  |
|  | B2 | 1.5±0.1 | 1.5 | B2 | 5.6±0.4 |  |
|  | B3 | 1.3±0.1 |  | B3 | 4.4±0.4 | 5.2 |
|  | B4 | 1.5±0.1 |  | B4 | 5.0±0.4 |  |
|  | B5 | 1.5±0.1 |  | B5 | 5.1±0.4 |  |
| WT | A1 | 2.6±0.1 |  | A1 | 4.3±0.6 |  |
|  | A2/A5 | 2.5±0.1 | 2.4 | A2/A5 | 5.6±0.8 | 4.5 |
|  | A3 | 2.0±0.1 |  | A3 | 3.7±0.5 |  |
|  | Ch1 | 1.9±0.1 |  | Ch1 | 6.1±0.5 |  |
|  | Ch2 | 1.8±0.1 | 2.1 | Ch2 | 4.6±0.5 | 5.4 |
|  | Ch4 | 2.8±0.2 |  | Ch4 | 5.6±0.7 |  |
|  | B1 | 1.9±0.1 |  | B1 | 6.7±0.7 |  |
|  | B2 | 1.6±0.1 |  | B2 | 6.2±0.6 |  |
| Δ*sglA* | B3 | 1.5±0.1 | 1.7 | B3 | 5.1±0.5 | 5.9 |
|  | B4 | 1.8±0.1 |  | B4 | 5.4±0.5 |  |
|  | B5 | 1.6±0.1 |  | B5 | 6.1±0.6 |  |
|  | A1 | 3.2±0.1 |  | A1 | 5.8±0.1 |  |
|  | A2/A5 | 3.2±0.2 | 3.0 | A2/A5 | 5.4±0.1 | 4.9 |
|  | A3 | 2.6±0.1 |  | A3 | 3.4±0.9 |  |
|  | Ch1 | 1.8±0.1 |  | Ch1 | 6.7±0.7 |  |
|  | Ch2 | 2.0±0.1 | 2.3 | Ch2 | 5.4±0.7 | 6.5 |
|  | Ch4 | 3.0±0.1 |  | Ch4 | 7.4±0.9 |  |

**Table S6**. **Recipe of mineral-based solid medium.** The pH is adjusted to 6.5 with H_3_PO_4_ or 0.25 M KOH. Each sample uses 100 mL of medium that contains 2g agar, 2 g of ^13^C-glucose and 5 mL of sodium nitrate solution with 0.1ml of trace elements. The culture media and condition were adapted from a previously described protocol^1^.

|  | Reagent | Quantity for 1L |
| --- | --- | --- |
| Trace elements | ZnSO_4_.7H_2_O | 22.0g |
|  | H_3_BO_3_ | 11.0g |
|  | MnCl_2_.4H_2_O | 5.0g |
|  | FeSO_4_.7H_2_O | 5.0g |
|  | CuSO_4_.5H_2_O | 1.6.0g |
|  | CoCl_2_.6H_2_O | 1.6.0g |
|  | (NH_4_)_6_Mo_7_O_24_.4H_2_O | 1.1.0g |
|  | EDTA | 50.0g |
| Nitrate salt solution | NaNO_3_ | 300.0g |
|  | KCl | 26.0g |
|  | MgSO_4_.7H_2_O | 24.0g |
|  | KH_2_PO_4_ | 120 |
|  | K_2_HPO_4_ | 20.9 |
| Carbon source | Glucose | 10.0 |

**Table S7.**  **^13^C chemical shifts of *A*. *fumigatus* conidial cell wall.** For each carbon site, the ^13^C chemical shifts are shown in the top and bottom rows, respectively. The referencing scale is TMS scale for ^13^C, with ambiguity are underlined:

| Carbohydrates | forms | C1 | C2 | C3 | C4 | C5 | C6 | Reference |
| --- | --- | --- | --- | --- | --- | --- | --- | --- |
| Rigid molecules | | | | | | | | |
| β -1,3-glucan (B) |  | 103.6 | 74.4 | 86.8 | 68.4 | 77.6 | 61.4 | Shim *et al*. 2007^2^ |
| α-1,3-glucan | a | 101.2 | 71.9 | 84.9 | 69.9 | 71.6 | 60.9 | Bhanja et al.2014^3^ |
|  | f | 101.6 | 70.8 | 85.1 | 69.7 | 71.9 | 60.6 |  |
| Chitosan |  | 102.2 | 55.9 | 73.0 | 80.4 | 75.7 | - |  |
| Chitin | a | 103.9 | 55.3 | 73.7 | 83.5 | 75.6 | 60.7 | Fernando *et al.* 2021^4^ |
|  | b | 103.2 | 54.4 | 73.4 | 82.9 | 75.6 | 60.8 |  |
|  | c | 104.5 | 55.0 | 73.5 | 83.5 | 75.6 |  |  |
| Mannan |  | 102.3 | 71 | 63.9 | 67.1 | 75.8 | 60.6 |  |
| B-1,6-glucan | **-** | - | - | 76.7 | 70.5 |  | 69.4 | Lowman *et al.* 2011 ^5^ |
| Mobile molecules | | | | | | | | |
| β-1,3-glucan (B) |  | 103.6 | 74.4 | 86.7 | 68.3 | 77.6 | 61.5 | Shim *et al.* 2007^6^  Fairweather *et al.* 2009^7^  Saito *et al.* 1979^8^ |
| β-1,5 galactofuranose (Gal*f*) | a | 108 | 81.8 | 77.5 | 83.5 | 71.7 | 63.5 | Chakraborty *et al.* 2021^9^ |
|  | b | 107.5 | 81.9 | 77.4 |  |  |  |  |
| α-1,6-Mannan (Mn^1,6^) |  | 102.7 | 70.8 | 73.8 | 67.7 | 70.8 | --- | Latge *et al.* 1994^10^  Chakraborty *et al.* 2021^9^ |
| α-1,2-Mannan (Mn^1,2^) | a | 101.1 | 78.5 | 70.9 | 67.8 | 74.0 | 62.1 |  |
|  | b | 98.9 | 79.0 | 71.2 | 67.0 | 73.4 | 62.0 |  |
| Galactose/Glucose  or their derivatives  (Gl) | a | 90.0 | 74.0 | 74.5 | 70.3 | --- | 61.8 | Fontaine *et al.* 2011^11^ |
|  | b | 97.0 | 71.1 | 74.4 | 67.0 | --- | 61.8 | Archbald et al. 1981^12^ |
|  | c | 94.7 | 72.5 | 73.6 | 67.4 | --- | 61.9 | Archbald et al. 1981^12^  Fontaine *et al.* 2011^11^ |

**Table S8. Solid-state NMR experiments and parameters.** To be quantitative, direct pulse (DP) experiments with 35 s long recycling delay were used. cross polarization (CP), most rigid molecules. With DP and a shorter recycling delay of 2 seconds, suppress the rigid molecules from the spectra, and with Insensitive Nuclei Enhanced by Polarization Transfer (INEPT) the most mobile molecules were selected. For 2D ^13^C-^13^C correlation experiments allowed to resolve rigid intramolecular peaks. 2D DQ-SQ, DP J-INADEQAUTE spectra was used to detect through-bond correlations. The experimental parameters include the ^1^H Larmor frequency, total experiment time (t), recycle delay (d1), number of scans (NS), The number of points for the direct (td2) and indirect (td1) dimensions, the acquisition time of the direct dimension (aq2) and the evolution time of indirect dimension (aq1), spectral width (sw1 and sw2).

| Experiment | B_0_  (T) | t  (h) | d1  (s) | NS | td2 | td1 | aq2 (ms) | aq1 (ms) | sw2 (ppm) | sw1 (ppm) |
| --- | --- | --- | --- | --- | --- | --- | --- | --- | --- | --- |
| 1D ^13^C CP | 18.8 | 0.5 | 1.8 | 1024 | 3600 |  | 18 |  | 496.8 |  |
| 1D ^13^C DP | 18.8 | 0.5 | 2.0 | 256 | 3600 |  | 18 |  | 496.8 |  |
| 1D ^13^C DP | 18.8 | 2.5 | 35.0 | 256 | 3600 |  | 18 |  | 496.8 |  |
| 1D ^13^C refocused INEPT | 18.8 | 1.5 | 4.0 | 1024 | 3200 |  | 16 |  | 496.8 |  |
| 2D ^13^C-^13^C with CORD mixing | 18.8 | 11 | 2.0 | 32 | 2800 | 600 | 14 | 7.5 | 496.8 | 198.7 |
| 2D ^13^C refocused DP J-INADEQUATE | 18.8 | 6 | 1.5 | 16 | 2600 | 1024 | 19 | 10 | 326.8 | 248.5 |
| 1D ^13^C T_1_ | 9.4 | 3.5 | 2.0 | 512 | 2000 |  | 16 |  | 623.3 |  |
| 1D ^1^H T_1ρ_ | 9.4 | 4.0 | 2.0 | 512 | 2000 |  | 16 |  | 623.3 |  |
| 2D ^13^C-^13^C water-edited | 9.4 | 8.0 | 2.0 | 64 | 2000 | 220 | 16 | 5.4 | 623.3 | 199.4 |

**Table S9. ^1^H and ^13^C chemical shifts of *A. fumigatus* mobile polysaccharides from ^1^H-detection experiments.** For each carbon site, the ^13^C and ^1^H chemical shifts are shown in the top and bottom rows, respectively. The referencing scale is TMS scale for ^13^C, and DSS for ^1^H. ^13^C and ^1^H sites with ambiguity are underlined for Gl, the carbon numbering of C2, 3, 4 is uncertain.

|  | forms | C1 | C2 | C3 | C4 | C5 | C6 | Reference |
| --- | --- | --- | --- | --- | --- | --- | --- | --- |
| Rigid molecules | | | | | | | | |
| β -1,3-glucan (B) | a | --- | --- | 85.34  3.95 | --- | --- | --- | Shim *et al.* 2007^2^ |
|  | b | 104.5  4.6 | 74.4  3.5 | 84.75  4.1 | 70.34  3.6 | - | 61.7  3.7,3.8 | Fairweather *et al.* 2009^7^ |
|  | c | --- | 74.5  3.5/4.3 | 84.75  4.25 | 70.86  --- | --- | 62.36  3.8,3.9 | Saito *et al.* 1979^8^ |
|  | d | --- | 74.3 | 86  4.19 | 71.05 | --- | 62.5  3.8,3.9 |  |
| α-1,6-Mannan (Mn^1,6^) | a | 102.8  5.04 | 71.00  4.07/3.45 | --- | 67.0  3.6/3.8 | 71.00  4.07/3.45 | 61.85  3.7/3.8 | Latge *et al.* 1994^13^ |
|  | b | 99.10  4.9 | 71.02  3.9 | 72.5  3.8/3.5 | 67.04  3.93 | 71.02  3.9 | 61.44  3.7/3.8 | Chakraborty *et al.* 2021^14^ |
|  | c | 100.25  4.9 | 70.2  3.9 | --- | 67.04  3.93 | 70.2  3.9 | 61.44  3.7/3.8 | Kuraoka *et al.* 2021^15^ |
| α-1,2-Mannan (Mn^1,2^) | a | 101.3  5.2 | 78.67  4.09 | 70.90  3.95 | 67.7  3.65 | 74.04 | 61.90  3.80/3.7 | Kuraoka *et al.* 2021^15^ |
|  | b | 101.3  5.2 | 78.67  4.09 | 70.9  3.95 | 67.70  3.65 | 74.04 | 61.9  3.88,3.70 | Kuraoka *et al.* 2018^16^ |
|  | c | 99.07  5.08 | 79.34  4.01 | 71.12  3.95 | 67.24  3.79 | --- | 61.79  3.60/ 3.80 |  |
|  | a | 90.00  5.88 | --- | 74.4  4.30 | 70.30  4.20 | --- | 61.70  3.88/ 3.78 | Fontaine *et al.* 2011^17^ |
|  | b | 96.95  5.45 |  | 70.91  3.9 | 67.17  3.73 | --- | 61.7  3.7/3.8 |  |
|  | c | 94.55  4.90 | 71.94  3.63 | --- | 67.40  3.5 | --- | 61.75  3.76 | Archbald et al. 1981^18^ |
| Galactose/Glucose or their derivatives (GI) | d | 94.85  5.17 | --- | --- | --- | --- | --- | Fontaine *et al.* 2011^17^ |
|  | e | 89.23  6.05 | --- | 74.85  4.12 | 71.3  4.40/3.6 | --- | 62.20  3.80, 3.90 |  |
|  | f | 94.77  4.94 | 71.87  3.92 | --- | 68.82  3.79 | --- | 63.85  3.7 |  |
|  | g | 88.64  5.87 | --- | 74.35  3.5 | 71.21  3.64 | --- | 62.25  3.85 |  |
| Glucose (Glc) | a (α) | 92.94  5.20 | 72.40  3.50 | 72.40  3.50 | 70.60  3.40 | --- | 61.56  3.70, 3.80 | Archbald *et al.* 1981^18^ |
|  | b (β) | b (β) | 96.84  4.63 | 74.90  3.25 | 76.60  3.50 | 70.50  3.40 | --- |  |
| GalNAc |  | 96.87  4.61 | 55.61  3.61 | 74.9  3.25 | 76.6  3.50 | 70.5  3.34 | 61.65  3.7, 3.88 |  |
| GalN |  | 92.94  5.22 | 54.94  3.69 | 72.4  3.5 | 70.9  3.4 | --- | 61.4  3.7/3.83 |  |
| Galf | a | 108.51  5.05 | 81.8  4.13 | 77.65  4.07 | 83.58  4.07 | 71.68  3.84 | 63.75  3.67/3.84 |  |
|  | b | 107.67  5.21 | 82.13  4.13 | 77.27  4.07 | 83.01  4.0 | 71.13  3.9 | 63.75  3.67/3.84 |  |

**Table S10. ^1^H and ^13^C chemical shifts of rigid polysaccharides from ^1^H-detection experiments.** For each carbon site, the ^13^C and ^1^H chemical shifts are shown in the top and bottom rows, respectively. The referencing scale is TMS scale for ^13^C, and DSS for ^1^H.

| Carbohydrates | C1 | C2 | C3 | C4 | C5 | C6 | Reference |
| --- | --- | --- | --- | --- | --- | --- | --- |
| β -1,3-glucan | 103.6  5.35 | 74.4  3.9 | 86.8  3.9 | 68.4  3.85 | 77.6  3.7 | 61.4  4.2 | Shim *et al*. 2007 |
| α-1,3-glucan  type-a | 101.1  5.7 | 71.9  3.5,3.8,  4.1,4.5 | 84.9  3.9 | 69.9  4.0 | 71.6  3.5, 3.8, 4.1, 4.5 | 60.9  4.15 | Bhanja et al.2014  Yarava *et al*. 2025 |
| α-1,3-glucan  type-d | 101.1  5.7 | --- | 84.41  4.40 | --- | --- | --- | Yarava *et al*. 2025 |
| α-1,3-glucan  type-e | 100.76  5.4 | --- | 84.38  3.57 | --- | --- | --- | Yarava *et al*. 2025 |
| α-1,3-glucan  type-f | 101.42  5.2 | --- | 84.38  3.57 | --- | --- | --- |  |
| α-1,3-glucan  type-g | 101.03  6.58 | --- | 84.37  4.81 | --- | --- | --- |  |
| α-1,3-glucan  type-h | 101.16.0 |  | 84.9  3.9 |  |  |  |  |
| Chitosan | 102.2  4.7, 4.3,  5.87 | 55.9  4.21 | 73.0  4.15 | 80.4  5.0 | 75.7  3.85 | - |  |
| Chitin type-a | 103.9  5.35 | 55.3  4.5 | 73.7  4.15 | 83.5  3.85 | 75.3  3.9 | 61.7  4.2 | Fernando *et al.* 2021 |

**Table S11**. Experimental parameters used for both WT and mutant A. fumigatus. Rigid region of the cell wall was characterized using CP based experiments on 600 MHz (14.1T) spectrometer using 1.3mm triple resonance MAS probe with the MAS frequency of 60 kHz.

| Expt. | CP (µs) | | NS | d1  (s) | td2 | td1 | td3 | aq2 (ms) | aq1 (ms) | aq3 (ms) | DIPSI-3 (ms) | RFDR mixing  (ms) | Expt.  Time (h) |
| --- | --- | --- | --- | --- | --- | --- | --- | --- | --- | --- | --- | --- | --- |
|  | t_cp1_ | t_cp2_ |  |  |  |  |  |  |  |  |  |  |  |
| 2D hCH | 500  (HC-CP) | 50  (CH-CP) | 32 | 2 | 1764  (^1^H) | 448  (^13^C) | - | 14.9 | 7.46 | - | - | - | 8h 34m |
| 2D hChH (RFDR) | 500  (HC-CP) | 500  (CH-CP) | 32 | 2 | 1764  (^1^H) | 448  (^13^C) | - | 14.9 | 7.46 | - | - | 0.533 | 8h  28m |

**Table S12**. **Experimental parameters used for both *A. fumigatus* WT and mutant samples.** Mobile region of the cell wall was characterized using scalar-coupling (*J*) based experiments on 800 MHz (18.8T) spectrometer using 3.2 mm triple resonance MAS probe with the MAS frequency of 15 kHz.

| Experiment | MAS | d1 | NS | td2 | td1 | td3 | aq2 (ms) | aq1 (ms) | aq3 (ms) | Decoupling/  Water suppression | *J-*evolution (ms) | DIPSI-3 (ms) | Expt.  Time (h) |
| --- | --- | --- | --- | --- | --- | --- | --- | --- | --- | --- | --- | --- | --- |
|  |  |  |  |  |  |  |  |  |  | SPINAL-64  (rf 71.429 kHz)  WALTZ-16  (rf 17 kHz)  MISSISSIPI  (total duration)  40 ms  (rf 25.994 kHz |  |  |  |
| 2D hCCH TOCSY  (DIPSI-3) | 15 | 2 | 32 | 2614  (^1^H) | 512  (^13^C) | 1 | 39.9 | 10.2 | - |  | 1.78 (τ_1_)  1.19 (τ_2_ | 25.5 | 9h 42m |
| 3D hCCH TOCSY  (DIPSI-3) | 15 | 2 | 8 | 2614  (^1^H) | 128  (^13^C) | 128  (^13^C) | 39.9 | 2.56 | 2.56 |  |  |  | 77h 53m |
